# Molecular Basis for CPC-Sgo1 Interaction: Implications for Centromere Localisation and Function of the CPC

**DOI:** 10.1101/2021.08.27.457910

**Authors:** Maria Alba Abad, Tanmay Gupta, Michael A Hadders, Amanda Meppelink, J Pepijn Wopken, Elizabeth Blackburn, Juan Zou, Lana Buzuk, David A Kelly, Toni McHugh, Juri Rappsilber, Susanne M A Lens, A Arockia Jeyaprakash

## Abstract

The Chromosomal Passenger Complex (CPC; consisting of Borealin, Survivin, INCENP and Aurora B kinase) and Shugoshin 1 (Sgo1) are key regulators of chromosome bi-orientation, a process essential for error-free chromosome segregation. Their functions rely on their ability to associate with centromeres. Two histone phosphorylations, histone H3 Thr3 (H3T3ph; directly recognised by Survivin) and histone H2A Thr120 (H2AT120ph; indirectly recognised via Sgo1), together with CPC’s intrinsic ability to bind nucleosome, facilitate CPC centromere recruitment. The molecular basis for CPC-Sgo1 binding and how their direct interaction influences CPC centromere localisation and function are lacking. Here, using an integrative structure-function approach, we show that the histone H3-like Sgo1 N-terminal tail interacts with Survivin acting as a hot-spot for CPC-Sgo1 assembly, while downstream Sgo1 residues, mainly with Borealin contributes for high affinity interaction. Disruption of the Sgo1 N-terminal tail-Survivin interaction abolished CPC-Sgo1 assembly *in vitro* and perturbed centromere localisation and function of CPC. Our findings provide evidence that CPC binding to Sgo1 and histone H3 N-terminal tail are mutually exclusive, suggesting that these interactions will likely take place in a spatially/temporally restricted manner and provide a rationale for the Sgo1-mediated ‘kinetochore proximal centromere’ pool of CPC.

## Introduction

Equal and identical segregation of chromosomes to the daughter cells during mitosis requires physical attachment of duplicated sister chromatids (via their kinetochores) to microtubules emanating from opposite spindle poles and subsequent alignment of chromosomes at the metaphase plate, a state known as bi-orientation (Musacchio and Desai, 2017). Chromosome bi-orientation is achieved and monitored by several processes including sister chromatid cohesion and quality control mechanisms known as error-correction and spindle assembly checkpoint (SAC), all controlled by the spatiotemporal regulation of kinases and phosphatases (Funabiki and Wynne, 2013; Gelens et al., 2018; Saurin, 2018).

Cohesin, a ring-shaped protein complex is a major player that mediates sister chromatid cohesion in S-phase (Haering et al., 2008; Haering et al., 2002). During prophase, the bulk of cohesin is removed from the chromosome arms (Gandhi et al., 2006; Kueng et al., 2006), while centromeric cohesin is maintained until anaphase onset protected by Shugoshin 1 (Sgo1) (Kitajima et al., 2006; Salic et al., 2004). Cdk1 phosphorylation of Sgo1 during mitosis enables the binding of the Sgo1-protein phosphatase 2 (PP2A) complex to cohesin and ensures that the two sister chromatids remain connected until anaphase onset, when separase cleaves the remaining centromeric cohesion allowing the sister chromatids to separate (Kitajima et al., 2006; Liu et al., 2013b; Shintomi and Hirano, 2009; Waizenegger et al., 2000). Sgo1 localisation to centromeres is crucial for its role as cohesion protector. Sgo1 has been suggested to first localise to kinetochores via the Bub1 dependent histone H2A phosphorylation at T120 (H2AT120ph) in order to then efficiently load onto centromeres to protect cohesion and prevent premature sister chromatid separation (Broad et al., 2020; Hengeveld et al., 2017; Kawashima et al., 2010; Liu et al., 2013a).

Error correction is a mechanism that destabilises incorrect kinetochore-microtubule (KT-MT) attachments, such as syntelic (two sister kinetochores attached to microtubules from the same spindle pole) or merotelic (single kinetochore attached to microtubules emanating from both spindle poles) attachments, and stabilises correct bi-polar attachments. The chromosomal passenger complex (CPC), consisting of Aurora B kinase, INCENP, Borealin and Survivin is one of the key players regulating this process (Carmena et al., 2012). The CPC, via its Aurora B enzymatic core, destabilises aberrant KT-MT attachments by phosphorylating outer kinetochore substrates such as the Knl1/Mis12 complex/Ndc80 complex (KMN) network so that new attachments can be formed (Cheeseman et al., 2006; Cimini et al., 2006; DeLuca et al., 2006; Lampson et al., 2004; Welburn et al., 2010). Sgo1 has also been shown to regulate KT-MT attachments via PP2A-B56 recruitment that balances Aurora B activity at the centromeres (Meppelink et al., 2015). In addition to error correction, the CPC is also involved in the regulation of the SAC, a surveillance mechanism that prevents anaphase onset until all kinetochores are attached to microtubules (Foley and Kapoor, 2013; Musacchio, 2015).

CPC function relies on its ability to localise correctly during mitosis. The CPC predominantly localises in the centromeric region between the sister-kinetochores during early mitosis and three independent studies recently suggested that the evolutionary conserved Haspin and Bub1 kinases can recruit independent pools of the CPC along the inter-kinetochore axis. Both recruitment pathways appear redundant for KT-MT error correction and can support faithful chromosome segregation (Broad et al., 2020; Hadders et al., 2020; Liang et al., 2019). Haspin mediates phosphorylation on histone H3 Thr3 (H3T3ph), which is recognised by the BIR domain of Survivin (Jeyaprakash et al., 2011; Kelly et al., 2010; Wang et al., 2010). Bub1 phosphorylates Thr120 of Histone H2A (H2AT120ph) (Bonner et al., 2020; Tsukahara et al., 2010; Wang et al., 2010; Yamagishi et al., 2010) that is recognised by Sgo1 which in turn, is suggested to interact with Borealin via its coiled-coil domain (Bonner et al., 2020; Yamagishi et al., 2010). However, our earlier work showed that Sgo1 N-terminal tail (AKER peptide) can also interact with Survivin BIR domain using a binding mode that is nearly identical to that of the histone H3 tail phosphorylated at Thr3 (Jeyaprakash et al., 2011), suggesting that a direct interaction between Survivin and Sgo1 could also be possible. H3T3ph and H2AT120ph appear to localise to distinct regions within the mitotic centromeres, with H3T3ph localising to the inner centromere and H2AT120ph to the KT-proximal outer centromere (Broad et al., 2020; Liu et al., 2013a; Yamagishi et al., 2010). While CPC-Sgo1 functional interdependency is well established, structural and molecular basis for how CPC and Sgo1 interact and the contribution of this physical interaction to the centromere localisation and function of the CPC remain unclear. Here, we address these questions by combining biochemical, structural, biophysical and cellular approaches.

## Results and discussion

### CPC-Sgo1 forms a robust complex *in vitro*

The CPC-Sgo1 interaction has been reported to be critical for sister chromatid bi-orientation and subsequent accurate chromosome segregation from yeast to humans (Hengeveld et al., 2017; Hindriksen et al., 2017; Peplowska et al., 2014; Tsukahara et al., 2010). However, the molecular basis of this interaction has not been characterised. To assess whether the CPC can directly interact with Sgo1 *in vitro*, we purified recombinant CPC containing INCENP_1-58_, full length Survivin and a stable version of Borealin lacking the first 9 residues, Borealin_10-280_ (CPC_ISB10-280_, Fig. 1A) and tested its interaction with recombinant Sgo1_1-415_ (just lacking the HP1 binding domain and the Sgo motif) using size exclusion chromatography (SEC) (Fig. 1A and S1A). Our data showed that Sgo1_1-415_ and CPC_ISB10-280_ can form a stable monodisperse complex *in vitro* as analysed by SEC (Fig. 1B). Using Isothermal Titration Calorimetry experiments (ITC) we assessed the binding affinity of this interaction. CPC_ISB10-280_ and Sgo1_1- 415_ exhibited high affinity with a K_d_ in the low nM range (K_d_= 53 ± 7 nM; Fig. 1C and S2C). The interaction is both enthalpically (ΔH= −6.58 ± 0.098 kcal.mol^-1^) and entropically (-TΔS= - 3.19 Kcal.mol^-1^) driven. ITC data also revealed a 1:1 stoichiometry for the CPC-Sgo1 complex, which fitted with the data obtained by mass photometry (Fig. S1B, S1C and S1D). Mass photometry analysis of the CPC_ISB10-280_/Sgo1_1-415_ complex showed a main population of 193 ± 29 kDa (Fig. S1D). Since the calculated MW for two molecules of Sgo1_1-415_ and two molecules of CPC_ISB10-280_ is 203.6 kDa, our data imply that a CPC_ISB10-280_ dimer (105 ± 17.5 kDa; Fig. S1B; calculated MW for a CPC_ISB10-280_ dimer is 108.8 kDa) is binding to a Sgo1_1-415_ dimer (82 ± 24 kDa; Fig. S1C; calculated MW for a Sgo1_1-415_ dimer is 94.8 kDa).

**Fig. 1.**
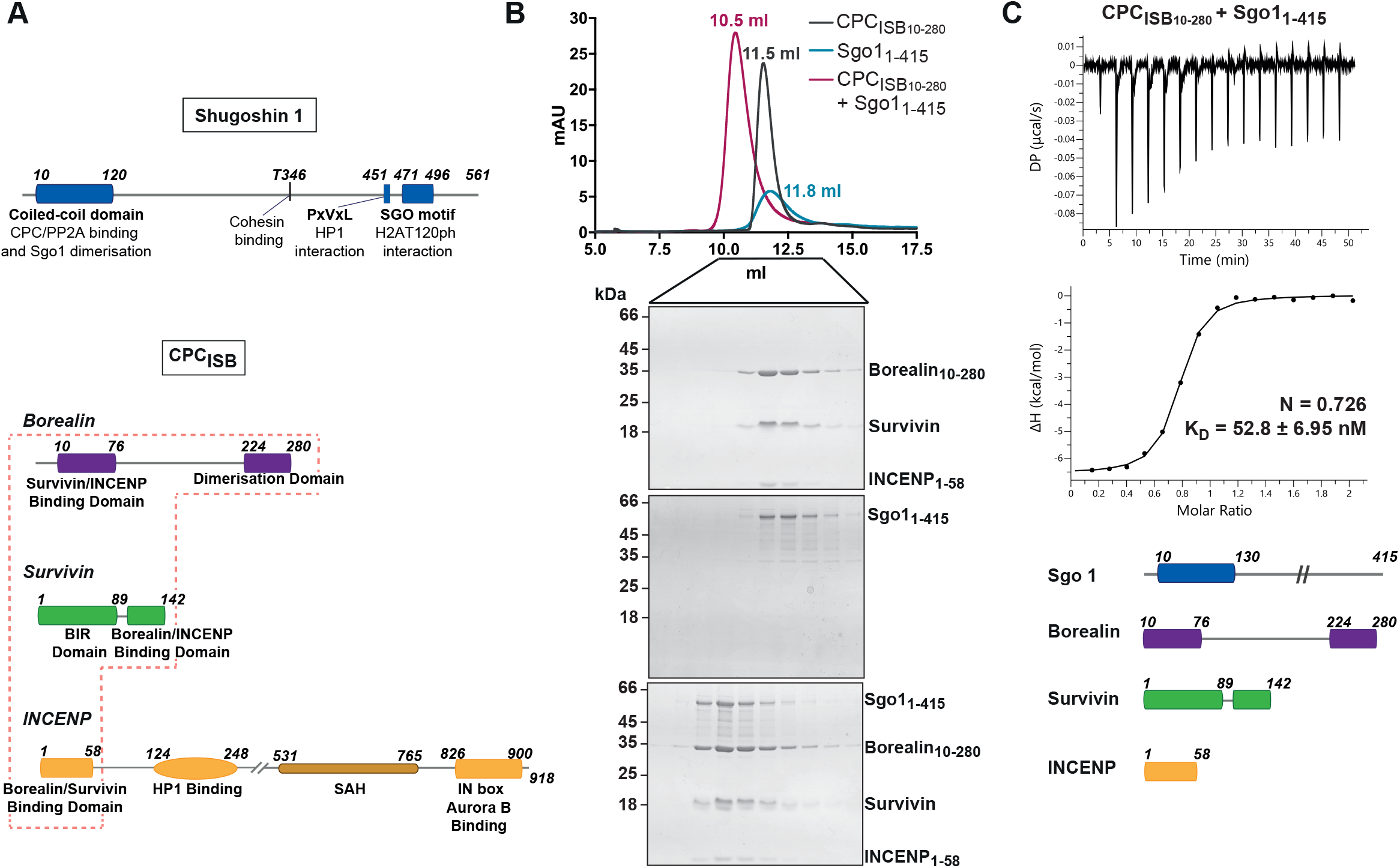
CPC-Sgo1 forms a robust complex *in vitro*. **A)** Schematic diagram depicting the domain architecture of Sgo1 and the Borealin, Survivin and INCENP subunits of the CPC. CPC_ISB_ (INCENP_1-58_, Survivin full length and Borealin full length) is highlighted in a red box. **B)** Size Exclusion Chromatography (SEC) profiles and corresponding Coomassie-stained SDS-PAGEs for the analysis of Sgo1_1-415_ (blue) and CPC_ISB10-280_ (INCENP_1-58_, Survivin and Borealin_10-280_; dark grey) complex formation (red). A Superdex S200 10/300 GL (Cytiva) column pre-equilibrated with 25 mM HEPES pH 8, 250 mM NaCl, 5 % Glycerol and 2 mM DTT was used. Elution volumes of peak fractions are indicated above the chromatogram peaks. **C)** Isotherms for Sgo1_1-415_ interaction with CPC_ISB10- 280_ (40 μl of 50 μM CPC_ISB10-280_ were injected into 200 μl of 5 μM Sgo1_1-415_). The ITC experiment was performed with 16 × 2.5 μl injections (0.5 μl 1^st^ injection), 180 sec apart at 20 °C. Top panel shows the raw ITC data, while the bottom panel shows the integrated heat data corrected for heat of dilution and fits to a standard 1:1 binding model (Malvern Instruments MicroCal Origin software, V1.3).

### Sgo1 makes multipartite interactions with CPC subunits

Previous studies have suggested that the Sgo1-CPC interaction is mediated via the N-terminal coiled-coil of Sgo1 and Borealin (Bonner et al., 2020; Tsukahara et al., 2010). However, our structural data revealed that the very N-terminus of Sgo1 can interact with the BIR domain of Survivin (Jeyaprakash et al., 2011). Together, these studies suggest that multipartite interactions between Sgo1 and different CPC subunits could facilitate CPC-Sgo1 complex formation. To gain further structural insights, we performed chemical cross-linking of the CPC_ISB10-280_-Sgo1_1-415_ complex using a zero-length cross-linker, 1-ethyl-3-(3-dimethylaminopropyl) carbodiimide (EDC), followed by mass spectrometry analysis (Fig. S1E). Cross-linking-mass spectrometry (CLMS) data showed that: (1) consistent with our previous observations (Jeyaprakash et al., 2011), the N-terminal region of Sgo1 (amino acids 1-34) makes extensive contacts with Survivin BIR domain (amino acids 18-89); (2) the N-terminal coiled-coil of Sgo1 (amino acids 10-120) interacts with the CPC triple helical bundle; (3) consistent with previous findings (Bonner et al., 2020), the N-terminal coiled-coil also contacts the Borealin dimerisation domain; and (4) the Sgo1 region beyond the N-terminal coil-coil region, which is predicted to be unstructured, contacts both Survivin and Borealin with most contacts confined to the Sgo1 central region spanning amino acids (aa) 180-300 (Fig. 2A and 2B). Thus, our cross-linking results suggests that Sgo1 interacts with CPC, mainly via two regions, the N-terminal coiled-coil domain and the unstructured central region (Fig. 2A and 2B).

**Fig. 2.**
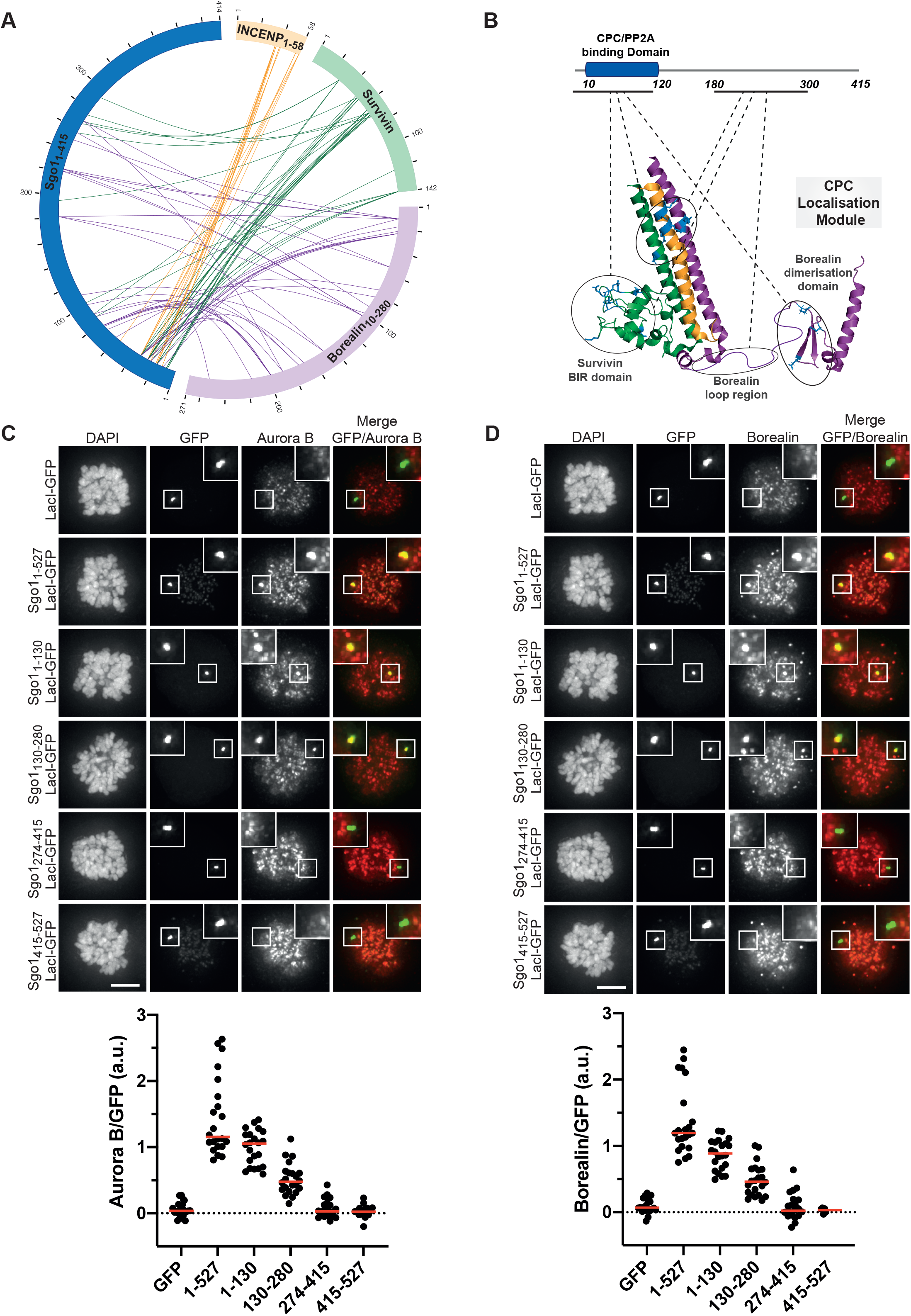
Sgo1 makes multipartite interactions with CPC components. **A)** Circular view of the EDC cross-links observed between the different subunits of the CPC_ISB10-280_ (INCENP_1-58_ in yellow, Survivin in green and Borealin_10-280_ in purple) and Sgo1_1-415_ (dark blue). For clarity, only contacts between Sgo1 and the CPC subunits are shown. Intermolecular contacts of INCENP, Survivin and Borealin with Sgo1 are shown as yellow, green and purple lines, respectively. XiNet (Kolbowski et al., 2018) was used for data visualisation. Auto-validation filter was used. **B)** Cartoon representation of the crystal/NMR structures of the CPC [CPC core PDB: 2QFA, (Jeyaprakash et al., 2007) and Borealin dimerisation domain PDB: 2KDD, (Bourhis et al., 2009)] and domain architecture of Sgo1 highlighting the regions involved in the CPC-Sgo1 contacts observed in A. **C and D)** Representative immunofluorescence images (top) and quantification (bottom) for the analysis of the recruitment of endogenous Aurora B (C) and Borealin (D) to the LacO array in U-2 OS-LacO Haspin CM cells expressing different Sgo1-LacI-GFP constructs (Sgo1_1-527_-LacI-GFP, Sgo1_1-130_-LacI-GFP, Sgo1_130-280_-LacI-GFP, Sgo1_274-415_-LacI-GFP, Sgo1_415-527_-LacI-GFP). The graphs show the intensities of Aurora B and Borealin over GFP (dots) and the means (red bar). Data are a representative of two independent experiments. Scale bar, 5 μm.

We further analysed the contribution of different Sgo1 regions for CPC binding using a LacO-LacI tethering assay. For this we made use of U-2 OS cells harbouring a LacO array on the short arm of chromosome 1 to which we could recruit Sgo1 fragments as LacI-GFP fusions (U-2 OS-LacO cells; Janicki et al., 2004). To exclude any contribution from H3T3ph on CPC recruitment we made use of a Haspin CRISPR mutant (CM) cell line that displays no discernible Haspin activity (Hadders et al., 2020). Constructs containing Sgo1_1-130_ and full length Sgo1_1-527_ recruited endogenous Aurora B (Fig. 2C) and Borealin (Fig. 2D) to the LacO foci, at comparable levels. This is in line with previous data that suggested the Sgo1 N-terminal region as a major CPC binding site (Bonner et al., 2020; Jeyaprakash et al., 2011; Tsukahara et al., 2010). Surprisingly, Sgo1_130-280_ fused to LacI-GFP was also able to recruit endogenous Aurora B (Fig. 2C) and Borealin (Fig. 2D) to the LacO foci, although at lower levels compared to Sgo1_1-130_ and Sgo1_1-527_. On the contrary, the Sgo1 fragments Sgo1_274-415_ and Sgo1_415-527_ (Sgo1_274-415_-LacI-GFP and Sgo1_415-527_-LacI-GFP) failed to recruit either Aurora B or Borealin (Fig. 2C and 2D). Taken together, our data confirm that the main CPC-interacting regions of Sgo1 lie within the N-terminal coiled-coil region of Sgo1 (Sgo1_1-130_) and the adjacent unstructured region (Sgo1_130-280_).

### The Survivin interaction with the Sgo1 N-terminal tail is essential for CPC-Sgo1 assembly

Our previous study identified a histone H3-like N-terminal tail in Sgo1 (Ala1-Lys2-Glu3-Arg4), which interacted with the Survivin BIR domain with similar affinity as did the histone H3 tail (Jeyaprakash et al., 2011). X-ray crystallographic structural analysis revealed that the mode of Sgo1 tail binding is near identical to that of the histone H3 tail with phosphorylated Threonine 3 (Ala1-Arg2-Thr3ph-Lys4; Jeyaprakash et al., 2011; Fig. S1F). However, whether the Sgo1 N-terminal tail interaction with Survivin is possible in the context of a longer Sgo1 fragment remained an open question. Here, using SEC, we confirmed that Sgo1 spanning aa residues 1-155 (Sgo1_1-155_; a shorter and stabler fragment identified from limited proteolysis of Sgo1_1-415_), can form a stable complex with Survivin (Fig. 3A), indicating that in the context of Sgo1_1-155_, the Sgo1 N-terminal tail is accessible for binding to Survivin.

**Fig. 3.**
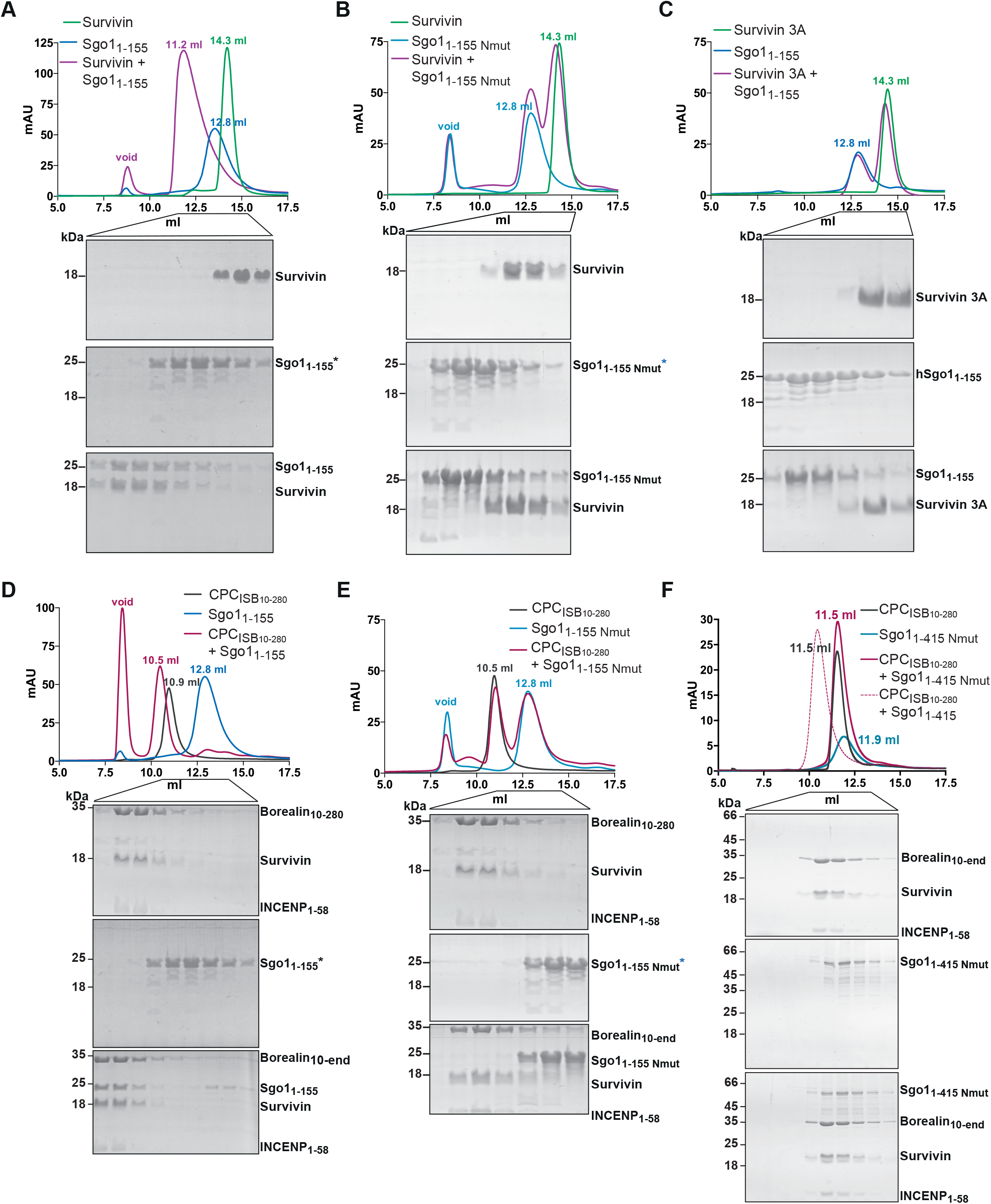
Survivin interaction with Sgo1 N-terminal tail is essential for CPC-Sgo1 assembly. **A-F)** Size Exclusion Chromatography profiles (top) and corresponding representative SDS-PAGEs stained with Coomassie (bottom) for the analysis of: **A)** Survivin and Sgo1_1-155_ interaction; **B)** Survivin and Sgo1_1-155 Nmut_ interaction; **C)** Survivin 3A and Sgo1_1-155_ interaction; **D)** CPC_ISB10-280_ and Sgo1_1-155_ interaction; **E)** CPC_ISB10-280_ and Sgo1_1-155 Nmut_ interaction and **F)** CPC_ISB10-280_ and Sgo1_1-415 Nmut_ interaction. A Superdex S200 10/300 GL (Cytiva) column pre-equilibrated with either 25 mM HEPES pH 7.5, 150 mM NaCl, 5 % Glycerol and 4 mM DTT (A-E) or 25 mM HEPES pH 8, 250 mM NaCl, 5 % Glycerol and 2 mM DTT (F) was used. Elution volumes are indicated on top of the chromatogram peaks.

Binding of histone H3 tail by Survivin requires anchoring of the small hydrophobic side chain of H3-Ala1 to the hydrophobic pocket of the Survivin BIR domain (Du et al., 2012; Jeyaprakash et al., 2011; Niedzialkowska et al., 2012). This interaction (anchoring of first Ala in the BIR pocket) is conserved in the Survivin-Sgo1 N-terminal peptide interaction (Jeyaprakash et al., 2011). To investigate the contribution of the Sgo1 N-terminal tail for binding to Survivin, we targeted this interaction by mutating the first alanine after the initiator methionine to a methionine (a residue with a long side chain not compatible with the BIR domain hydrophobic pocket; Sgo1_Nmut_). Remarkably, the Sgo1_1-155 Nmut_ mutant was unable to interact with Survivin, indicating that the Sgo1 N-terminal tail interaction with BIR domain is crucial for Survivin binding (Fig. 3B and S1F). Similarly, a Survivin BIR mutant (Survivin 3A: K62/E65/H80A) not capable of interacting with the histone H3 tail (Fig. S1F), failed to interact with Sgo1_1-155_ (Fig. 3C). These data together show that both the Sgo1 and histone H3 N-terminal tail use the same binding pocket in the Survivin BIR domain.

Consistent with our data (Fig. 1B), the shorter Sgo1_1-155_ fragment can also form a complex with CPC_ISB10-280_ *in vitro* (Fig. 3D). To evaluate the contribution of the Sgo1-Survivin interaction in the context of the CPC where Borealin can still interact with Sgo1 (Bonner et al., 2020; Tsukahara et al., 2010 and Fig. 2A and 2B), we mixed CPC_ISB10-280_ with Sgo1_1-155 Nmut_ (Fig. 3E) and Sgo1_1-415 Nmut_ (Fig. 3F) and tested their interaction by SEC. Strikingly, neither Sgo1_1- 155 Nmut_, nor the longer Sgo1_1-415 Nmut_ that includes the second CPC interacting region (aa 130-280) were able to interact with the CPC. This data agrees with the tethering assays where Sgo1_1- 130 Nmut_-LacI-GFP showed a drastic reduction in its ability to recruit Aurora B (Fig. S2A) and Borealin (Fig. S2B) to the LacO array as compared to Sgo1_1-130_-LacI-GFP. Together, our results reveal that the Survivin-Sgo1 interaction is essential for the CPC binding to Sgo1 and that the Sgo1 N-terminal tail acts as a hot-spot, whose perturbation abolishes the ability of CPC to form a complex with Sgo1.

### Borealin and INCENP are required for a high affinity CPC-Sgo1 interaction

To assess how different CPC subunits contribute to achieve high affinity Sgo1 interaction, we performed a series of Isothermal Calorimetry (ITC) experiments with either Survivin on its own or with CPC_ISB_ containing different Borealin truncations. Sgo1_1-130_ interacted with Survivin with mid-nanomolar affinity (K_d_ of 255 ± 33 nM; Fig. 4A and S2C). This, together with our previous observation that a Sgo1 N-terminal tail peptide bound Survivin with ∼ 1 μm affinity (Jeyaprakash et al., 2011) suggests that, although the interaction between the alanine and the Survivin BIR domain is essential for Sgo1/Survivin complex formation, the Sgo1-Survivin interaction extends beyond Sgo1 N-terminal tail. Sgo1_1-130_ bound CPC_ISB10-280_ with a K_d_ of 57 ± 8 nM, a ∼ 5 fold higher affinity as compared to the affinity for Survivin alone (Fig. 4B and S2C). This observation together with the CLMS analysis suggests that further interactions involving Borealin, and possibly INCENP, strengthen the affinity between the CPC and Sgo1. Consistent with our CLMS analysis (Fig. 2A and 2B) and a previous study (Bonner et al., 2020), CPC_ISB_ lacking the Borealin dimerisation domain (CPC_ISB10-221_) bound Sgo1_1-130_ with a three-fold lower affinity as compared to the CPC_ISB10-280_ (K_d_ =163 nM ± 16 nM vs 57 ± 8 nM), highlighting the contribution of the Borealin dimerisation domain for binding to Sgo1 (Fig. 4C and S2C). The measured affinity of CPC _ISB10-280_ binding to the near full length Sgo1 (Sgo1_1-415,_ K_d_ = 53 nM ± 7 nM, Fig. 1C), is almost identical to that for Sgo1_1- 130_ (Fig. 4B; K_d_ = 57 nM ± 8 nM). This confirms that the first 130 amino acids of Sgo1 represent the main CPC-interacting region *in vitro*. Furthermore, the observation that the affinity goes from a micromolar range for the AKER peptide with Survivin (Jeyaprakash et al., 2011) to the low nanomolar range for the CPC_ISB10-280_-Sgo1_1-130_ complex, indicates that although the interaction between the CPC and Sgo1 depends on the Sgo1 N-terminal tail binding to Survivin, the high-affinity interaction requires Sgo1 binding to Borealin and possibly INCENP. Overall, the ITC data indicates that the interaction between the Sgo1 N-terminal tail and Survivin is electrostatically driven (Fig. S2D), while the high-affinity interaction between the rest of the Sgo1 regions and the CPC is driven by entropic contribution that could be due to a release of water molecules associated with the surface and/or a conformational rearrangement upon binding (Fig. S2C). All these data together suggest that a weak micromolar affinity long-range electrostatic interaction between Survivin and the Sgo1 N-terminal tail is required to establish a high-affinity CPC-Sgo1 interaction mediated by multiple inter-protein contacts and hydrophobic effects.

**Fig. 4.**
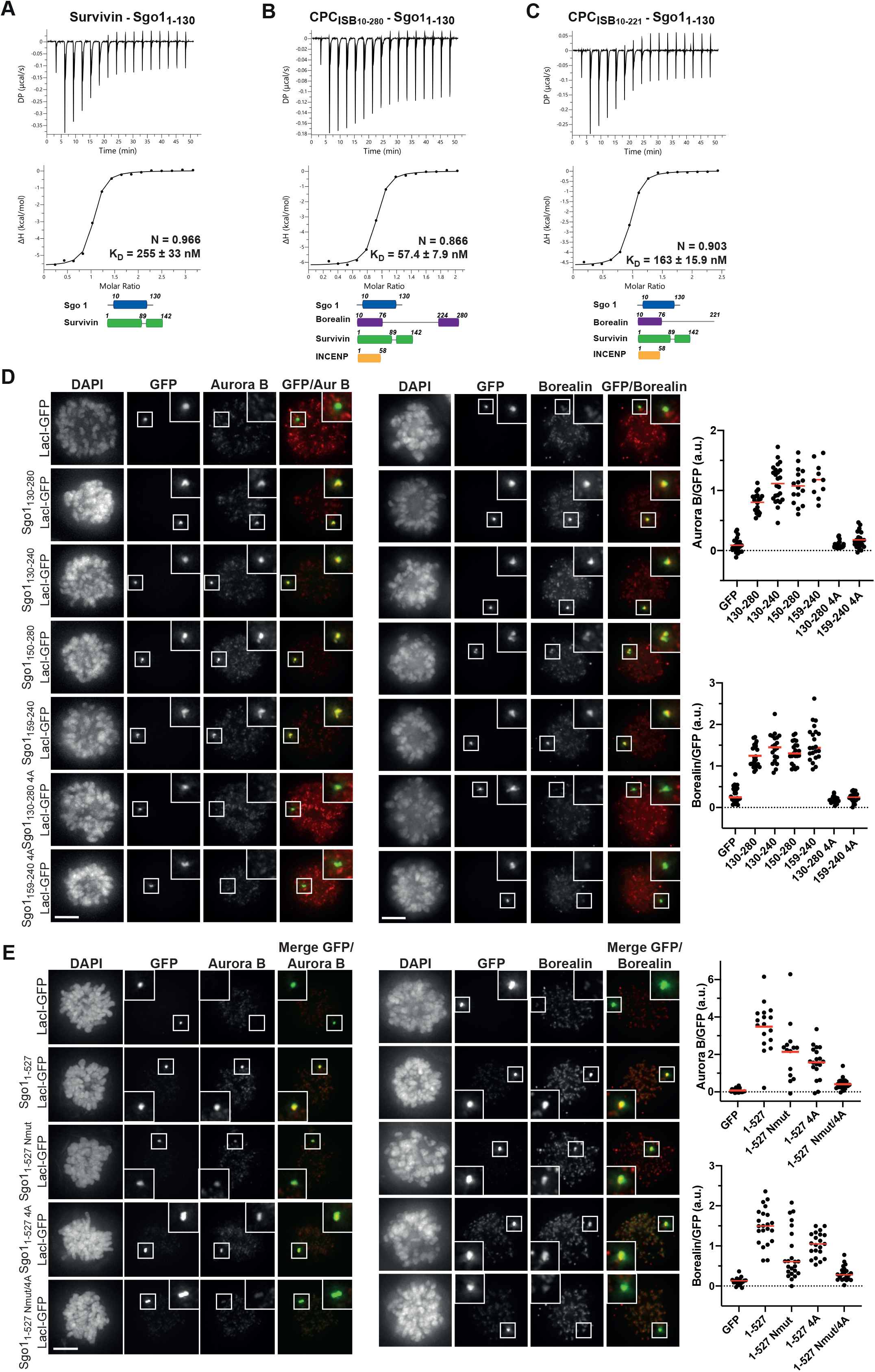
Multivalent interactions between Sgo1 and different CPC components are essential for high affinity CPC-Sgo1 binding and efficient CPC recruitment. **A-C)** Isotherms for the analyses of: **A)** Survivin interaction with Sgo1_1-130_ (40 μl of 312 μM Survivin was injected into 200 μl of 20 μM Sgo1_1-130_). **B)** CPC_ISB10-280_ interaction with Sgo1_1- 130_ (40 μl of 100 μM CPC_ISB10-280_ was injected into 200 μl of 10 μM Sgo1_1-130_). **C)** CPC_ISB10- 221_ interaction with Sgo1_1-130_ (40 μl of 240 μM CPC_ISB10-221_ was injected into 200 μl of 20 μM Sgo1_1-130_). The ITC experiments were performed with 16 × 2.5 μl injections (0.5 μl 1^st^ injection), 180 sec apart at 20 °C. Top panels show raw ITC data while bottom panels show integrated heat data corrected for heat of dilution and fit to a standard 1:1 binding model (Malvern Instruments MicroCal Origin software, V1.3). **D, E)** Representative micrographs (left) and quantifications (right) for the analysis of endogenous Aurora B and Borealin recruitment to the LacO array in U-2 OS-LacO Haspin CM cells expressing different Sgo1-LacI-GFP constructs: **D)** Sgo1_130-280_-LacI-GFP, Sgo1_130-240_-LacI-GFP, Sgo1_150-280_-LacI-GFP, Sgo1_159-240_-LacI-GFP, Sgo1_130-280 4A_-LacI-GFP or Sgo1_159-240 4A_-LacI-GFP; **E)** Sgo1_1-527_-LacI-GFP, Sgo1_1-527 Nmut_-LacI-GFP, Sgo1_1-527 4A_-LacI-GFP or Sgo1_1-527 Nmut/4A_-LacI-GFP. The graphs show the intensities of Aurora B and Borealin over GFP (dots) and the means (red bar). Data are a representative of four independent experiments. Scale bar, 5 μm.

### Interactions involving the Sgo1 N-terminal tail and a hydrophobic stretch spanning residues 188-191 are required for efficient recruitment of the CPC

Our cross-linking and tethering data (Fig. 2) identified an additional novel CPC-interacting region of Sgo1 within aa 130-280, which is found upstream of the cohesin binding site (Fig. 1A). To assess whether Sgo1_130-280_ can form a complex with the CPC *in vitro*, a 1.5 × molar excess of recombinant Sgo1_130-280_ was mixed with CPC_ISB10-280_ and the mix was analysed by SEC (Fig. S3A). SEC profiles and the analysis of SEC fractions showed that Sgo1_130-280_ can indeed form a complex with CPC_ISB10-280_. To further pinpoint the region within Sgo1_130-280_ that is necessary for the interaction with the CPC, we expressed Sgo1_130-280_ -LacI-GFP and multiple truncations of the 130-280 fragment in U-2 OS-LacO cells and assessed CPC recruitment through immunofluorescence analysis (Fig. 4D). A smaller fragment spanning Sgo1 amino acids 159-240 was capable of recruiting similar levels of the CPC as Sgo1_130-280_ -LacI-GFP (Fig. 4D). The region between 159-240 contained a highly conserved stretch of hydrophobic amino acids (188-191) and mutation of these residues to alanines (V188/S189/V190/R191A: 4A; Sgo1_130-280 4A_-LacI-GFP, Sgo1_159-240 4A_-LacI-GFP; Fig. S1A) completely abrogated CPC recruitment to both Sgo1_130-280_ and Sgo1_159-240_ to the CPC (Fig. 4D). Interestingly, when we introduced the same 4A mutation in recombinant Sgo1_130-280_, Sgo1_130-280 4A_, it still managed to interact with CPC_ISB10-280_ in the SEC analysis (Fig. S3B). This suggests that post-translational modifications within this Sgo1 region might facilitate its interaction with the CPC in cells.

As our analysis identified two CPC-interacting regions within Sgo1 (the N-terminal 130 aa including the N-terminal tail and the conserved coiled-coil, and the conserved hydrophobic region between aa 188 and 191), we next evaluated their contribution for CPC recruitment in the context of full length Sgo1 using the LacO tethering assay. Consistent with our *in vitro* data, full length Sgo1, harbouring the N-terminal mutation (Sgo1_1-527 Nmut_-LacI-GFP), recruited less Aurora B or Borealin (Fig. 4E) as compared with the Sgo1_1-527_-LacI-GFP. Similarly, the 4A mutation in the full length context (Sgo1_1-527 4A_-LacI-GFP) also reduced the recruitment of Aurora B and Borealin, while the double mutant (Sgo1_1-527 Nmut/4A_-LacI-GFP) showed an even stronger reduction of endogenous Aurora B and Borealin recruitment to the LacO array (Fig. 4E). Collectively, this data demonstrates the contribution of both Sgo1 regions for CPC binding in cells.

### The Survivin interaction with the Sgo1 N-terminal tail is essential for the centromeric localisation of the CPC and proper chromosome segregation

We next evaluated how the different Sgo1 regions we identified as important for the CPC-Sgo1 interaction contribute to maintaining the centromeric levels of the CPC in cells. Endogenous Sgo1 was depleted by siRNA in HeLa Kyoto cells expressing either wild type Sgo1 (Sgo1-GFP) or mutant Sgo1 (Sgo1_Nmut_-GFP, Sgo1_4A_-GFP or Sgo1_Nmut/4A_ double mutant) and centromeric levels of Borealin were analysed by quantitative immunofluorescence microscopy (Fig. 5A, S3C and S3D). Consistent with previous observations (Broad et al., 2020; Kawashima et al., 2007; Meppelink et al., 2015; Tsukahara et al., 2010; van der Waal et al., 2012; Wang et al., 2010), depletion of Sgo1 led to a two-fold reduction in the centromeric levels of Borealin. As expected, expression of wild type Sgo1 (Sgo1-GFP) rescued the centromeric abundance of Borealin (Fig. 5A). In line with our *in vitro* binding studies and cellular tethering data, expression of Sgo1 mutants (Sgo1_Nmut_-GFP, Sgo1_4A_-GFP or Sgo1_Nmut/4A_-GFP, Fig. S3D) aimed to perturb either the Sgo1 N-terminal tail-Survivin interaction or the Sgo1 188-191-Borealin interaction did not rescue the centromeric levels of Borealin, demonstrating direct contributions of these regions for efficient CPC recruitment to centromeres (Fig. 5A).

**Fig. 5.**
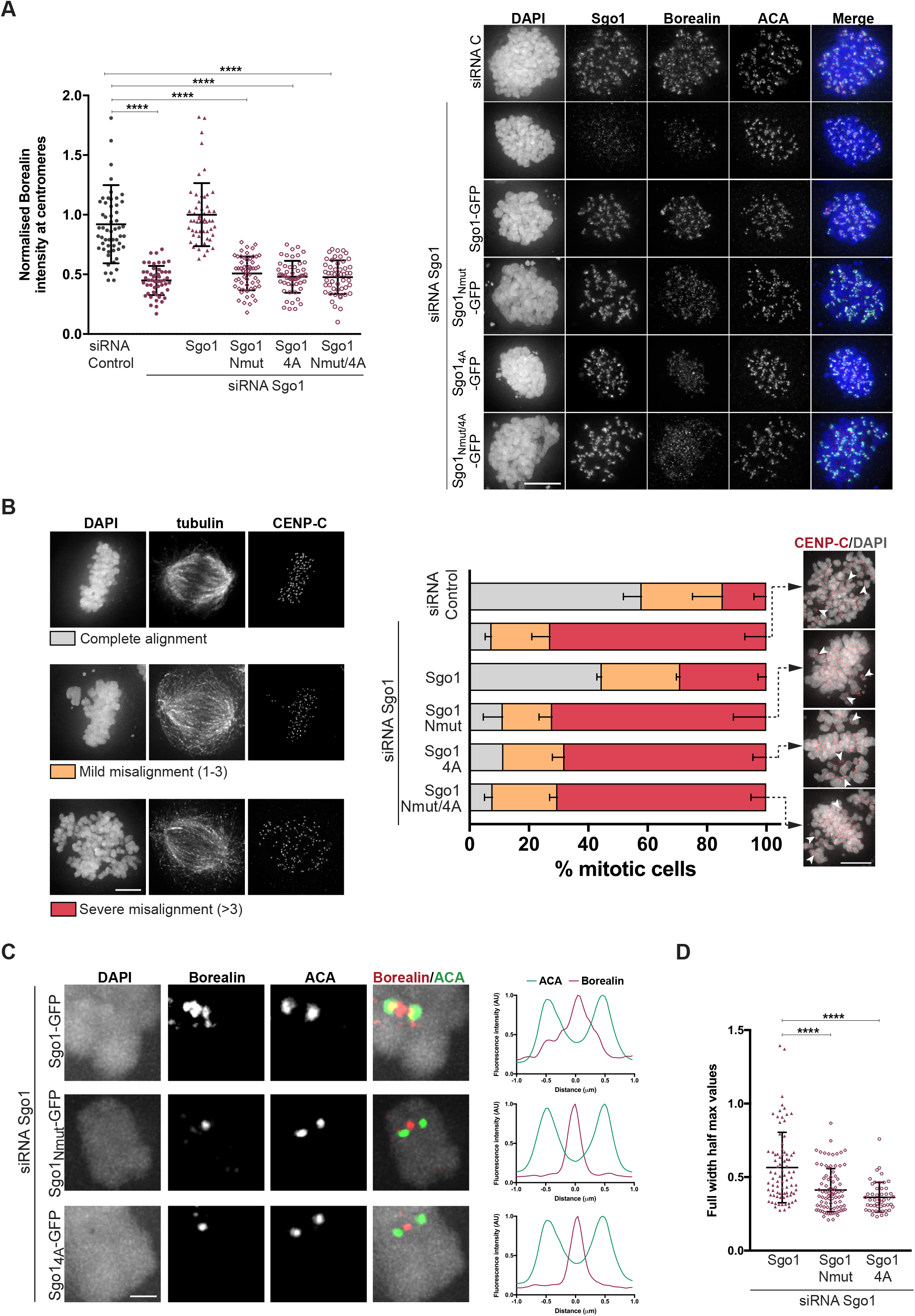
CPC interaction with the Sgo1 N-terminal tail is essential for the centromeric localisation of CPC and proper chromosome segregation. **A)** Representative micrographs of HeLa Kyoto cells expressing different Sgo1-GFP constructs (Sgo1-GFP, Sgo1_Nmut_-GFP, Sgo1_4A_-GFP or Sgo1_Nmut/4A_-GFP) and depleted of endogenous Sgo1 using siRNA oligos (right panel). Immunofluorescence of endogenous Borealin and ACA. DAPI was used for DNA staining. Scale bar, 10 µm. Quantification of Borealin levels at the centromeres using ACA as reference channel (left panel). Values normalised to Sgo1 siRNA/Sgo1-GFP condition. Three independent experiments, n ≥ 50, mean ± SD, Mann– Whitney test; ****, P ≤ 0.0001). **B)** Quantification of chromosome alignment of cells subjected to bi-orientation assay. Transfected cells were treated with 100 μM Monastrol for 16 h and released into medium containing 5 μM MG132 for 1 h. Representative examples of the alignment categories: complete alignment, mild misalignment (with 1-3 misaligned chromosomes) and severe misalignment (with more than 3 misaligned chromosomes) are found in the left panel. Representative images of the conditions expressing the three Sgo1 mutants showing pairs of CENP-C foci (red) (right panel). DAPI was used to visualise DNA. Scale bar, 5 μm. Three independent experiments, n ≥ 100; mean ± SD. **C)** Line plots depicting normalised fluorescence intensity levels of Borealin and ACA, measured along a line across the two sister ACA signals of the inter-kinetochore axis. Scale bar, 2 μm. Left, representative images of kinetochore pairs represented in the line plots. **D)** Quantification of the full width half max for the Borealin signal obtained in the line plots. Three independent experiments, n ≥ 49 kinetochores, mean ± SD, Mann–Whitney test; ****, P ≤ 0.0001).

Following the same experimental set-up described above, we depleted endogenous Sgo1 using siRNA oligos in HeLa Kyoto cells expressing wild type Sgo1 (Sgo1-GFP) or mutant Sgo1 (Sgo1_Nmut_-GFP, Sgo1_4A_-GFP or Sgo1_Nmut/4_-GFP) and quantified anaphase cells showing lagging chromosomes or chromosomes bridges, which are direct indicators of chromosome mis-segregation (Fig. S3E). This analysis confirmed that cells expressing Sgo1_Nmut_-GFP, Sgo1_4A_-GFP or Sgo1_Nmut/4A_-GFP show a high percentage of cells with either lagging chromosomes or chromosome bridges (27.8 ± 5.8 %, 26.9 ± 3.1 % and 29.3 ± 4.4 %, respectively) compared to the siRNA C and Sgo1-GFP rescue (6.1 ± 1.1 % and 10.3 ± 2.6 %, respectively).

We further analysed the effects of disrupting the CPC-Sgo1 interaction on chromosome bi-orientation. Sgo1-depleted HeLa cells expressing either Sgo1-GFP or mutant Sgo1 (Sgo1_Nmut_- GFP, Sgo1_4A_-GFP or Sgo1_Nmut/4_-GFP) were released from a monastrol-induced mitotic arrest into medium with MG132 and chromosome alignment was assessed. Expression of Sgo1 mutants led to around 70 % of cells with severe chromosome misalignment, comparable to the phenotype observed for Sgo1 depletion (Fig. 5B). Unlike Sgo1 knock-down cells, Sgo1-mutant expressing cells did not seem to experience loss of sister chromatid cohesion because sister centromeres remained closely together (Fig. 5B). This suggests that the alignment errors observed are not due to a loss of centromeric cohesion, but a reflection of perturbed kinetochore-microtubule error correction, presumably as a consequence of the reduced centromeric levels of CPC (Fig. 5A).

Considering the ability of Sgo1 to bind H2AT120ph and to recruit CPC to the kinetochore proximal centromere, we analysed the precise localisation of Borealin using chromosomes spreads of nocodazole-arrested HeLa cells expressing the Sgo1 mutants. Sgo1 depletion on Sgo1-GFP expressing HeLa cells showed Borealin mainly localised at the inner centromere but also displayed a small pool localised at the kinetochore proximal centromere (Fig. 5C), consistent with the previously described pattern of CPC localisation in unperturbed mitotic cells (Bekier et al., 2015; Hadders et al., 2020; Liang et al., 2019). On the contrary, depletion of Sgo1 on Sgo1_Nmut_-GFP or Sgo1_4A_-GFP expressing cells, Borealin was enriched as a single focus between the two sister ACA dots, similar to the inner centromere localisation previously observed for Borealin dimerisation mutants that bind less well to Sgo1 (Bekier et al., 2015). Quantification of the full width half max values for the Borealin intensity profiles obtained from the line plots of the chromosome spreads, confirmed that rescue of Sgo1 depletion with Sgo1-GFP expression generated a broader Borealin signal at the centromere (most likely the result of the combination of inner centromere and kinetochore proximal outer centromere pools), while expression of Sgo1 mutants (Sgo1_Nmut_-GFP and Sgo1_4A_-GFP) generated narrower Borealin profiles consistent with CPC localised at the inner centromere only. This data reveals that the interaction of CPC with H2AT120ph-bound Sgo1 is responsible for the kinetochore proximal centromere pool of the CPC.

A multi-pronged approach combining detailed biochemistry, CLMS analysis and available structural data with cellular phenotypic analysis provided us with an excellent opportunity to dissect the contributions of different Sgo1 regions and CPC subunits for CPC-Sgo1 complex formation. We show that both the Sgo1 N-terminal tail and the downstream hydrophobic region comprising aa 188-191 contribute to CPC-Sgo1 complex formation and are essential for efficient CPC centromere recruitment and function. Interestingly, the observation that the Sgo1 and histone H3 N-terminal tails exploit the same binding site in Survivin, suggests that these interactions are mutually exclusive and may explain why the Bub1-dependent CPC pool exists as a kinetochore-proximal centromere pool that is spatially distinct from the Haspin-dependent inner centromere CPC pool (Broad et al., 2020; Hadders et al., 2020; Liang et al., 2019). Future studies focusing on Sgo1 and CPC dynamics might provide further insights. It would also be interesting to see if and how CPC-Sgo1 interactions are modulated by CPC’s intrinsic ability to bind unmodified (Abad et al., 2019) and H3-T3ph nucleosomes and Sgo1 ability to bind H2AT120ph nucleosomes.

## Figure legends

**Fig. S1**.

**A)** Sequence alignment of Sgo1 orthologues from *Homo sapiens* (hs), *Bos taurus* (bt), *Mus musculus* (mm), *Gallus gallus* (gg), *Danio rerio* (dr) and *Xenopus laevis* (xl). The conservation score is mapped from red (highly conserved) to yellow (poorly conserved). Predicted secondary structure elements are shown below the sequence alignment. Multiple sequence alignment was performed with Clustal Omega (EMBL-EBI) and edited with Jalview 2.11.0 (Waterhouse et al., 2009). Highlighted with boxes are the N-terminal coiled-coil domain of Sgo1, the highly conserved 188-191 region and the Sgo motif. **B, C and D)** Resulting mass photometry histograms and kernel density estimates for CPC_ISB10-280_ (B), Sgo1_1-415_ (C) and CPC_ISB10-280_ /Sgo1_1-415_ complex. All samples were cross-linked with 0.01 % glutaraldehyde for 5 min at 4 °C. Mean ± SD. **E)** Representative SDS-PAGE analysis of CPC_ISB10-280_ cross-linked with Sgo1_1-415_ using EDC chemical cross-linker. **F)** Close up of the crystal structure of Survivin bound to a peptide comprising the four first amino acid residues of Sgo1 [AKER peptide; PDB: 4A0I, (Jeyaprakash et al., 2011)]. Sgo1_Nmut_ disrupts the interaction between the first amino acid of Sgo1 and the shallow hydrophobic pocket of Survivin. Mutation of amino acids Lys62, Glu65 and His80 in the Survivin BIR domain to alanine disrupt the crucial interactions with the AKER N-terminal tail of Sgo1.

**Fig. S2**.

**A, B)** Representative micrographs (top) and quantifications (bottom) for the analysis of endogenous Aurora B (A) and Borealin (B) recruitment to the LacO array in U-2 OS-LacO Haspin CM transfected with different Sgo1-LacI-GFP constructs: Sgo1_1-130_-LacI-GFP or Sgo1_1-130 Nmut_-LacI-GFP. The graphs show the intensities of Aurora B and Borealin over GFP (dots) and the means (red bar). Data are a representative of five independent experiments. Scale bar, 5 μm. **C)** Table including the ITC thermodynamic parameters for the different ITC experiments. **D)** Isotherms for the analyses of Survivin interaction with Sgo1_AKER_ peptide (40 μl of 375 μM Survivin was injected into 200 μl of 20 μM Sgo1_AKER_). The ITC experiment was performed with 16 × 2.5 μl injections (0.5 μl 1^st^ injection), 180 sec apart at 10 °C. Left panel shows raw ITC data while right panel shows integrated heat data corrected for heat of ligand dilution and fit to a standard 1:1 binding model (Malvern Instruments MicroCal Origin software, V1.3).

**Fig. S3**.

**A, B)** Size Exclusion Chromatography profiles (top) and corresponding representative SDS-PAGEs stained with Coomassie (bottom) for the analysis of CPC_ISB10-280_ and Sgo1_130-280_ interaction (A) and CPC_ISB10-280_ and Sgo1_130-280 4A_ (B) interaction. A Superdex S200 10/300 GL (Cytiva) column pre-equilibrated with 25 mM HEPES pH 8, 150 mM NaCl, 5 % Glycerol and 1 mM DTT was used. Elution volumes are indicated on top of the chromatogram peaks. **C)** Representative immunoblot for the analysis of Sgo1 levels upon Sgo1 depletion using siRNA oligos. Quantification of Sgo1/Tubulin ratio using uncalibrated OD values. Three independent experiments; mean ± SD; unpaired *t* test; ****, P ≤ 0.0001. **D)** Representative immunoblot of Sgo1-GFP constructs (Sgo1-GFP, Sgo1_Nmut_-GFP, Sgo1_4A_-GFP or Sgo1_Nmut/4A_-GFP) showing comparable expression levels. **E)** Quantification of anaphase cells with lagging chromosomes or chromosome bridges for the siRNA-rescue assay of the Sgo1-GFP constructs: Sgo1-GFP, Sgo1_Nmut-GFP_, Sgo1_4A-GFP_ or Sgo1_Nmut/4A-GFP_. Right panel shows representative examples of lagging chromosomes and chromosome bridges quantified. Three independent experiments, n ≥ 300; mean ± SD; unpaired *t* test; ns, not significant; **, P ≤ 0.01; ***, P ≤ 0.001).

## Materials and Methods

### Protein expression and purification of CPC and Sgo1

CPC_ISB10-280_ and CPC_ISB10-221_ were purified as previously described (Abad et al., 2019). Briefly, pRSET-His-GFP-3C-Survivin (pRSET vector from Thermo Fisher Scientific), pMCNcs-INCENP_1-58_ and pETM-His-TEV-Borealin_10-280_ or pETM-His-TEV-Borealin_10-221_ (pETM vector, gift from C. Romier, Institute of Genetics and Molecular and Cell Biology, Strasbourg, France) were co-transformed in BL21(DE3) pLysS. Cultures were grown at 37 °C until OD 0.8 and induced overnight at 18 °C with 0.35 mM IPTG. Cells were lysed in lysis buffer (25 mM HEPES pH 7.5, 500 mM NaCl, 25 mM Imidazole, 2 mM β-mercaptoethanol) and supplemented with complete EDTA-free cocktail tablets (Roche), 0.01 mg/ml DNase (Sigma) and 1 mM PMSF. The lysate was sonicated for 8 minutes, centrifuged at 58,000 × g for 50 minutes at 4°C and the complex was purified by affinity chromatography using His Trap Column (Cytiva). The protein-bound column was washed with lysis buffer followed by a high salt buffer wash (25 mM HEPES pH 7.5, 1 M NaCl, 25 mM Imidazole, 2 mM β-mercaptoethanol, 1mM ATP). The complex was eluted using high imidazole buffer (25 mM HEPES pH 7.5, 500 mM NaCl, 400 mM Imidazole, 2 mM β-mercaptoethanol) and the affinity tags were cleaved using TEV and 3C proteases while dialysing the sample in a buffer containing 25 mM HEPES pH 7.5, 150 mM NaCl and 4 mM DTT at 4° C overnight. The dialysed sample was then loaded onto a HiTrap SP HP (Cytiva) cation exchange column to separate the excess Borealin-Survivin complex and GFP from the CPC_ISB_ complexes. The samples containing stoichiometric and pure CPC_ISB_ complex were pooled, concentrated and run on a Superdex 200 Increase 10/300 (Cytiva) pre-equilibrated with 25 mM HEPES pH 8, 150 mM NaCl, 5 % Glycerol and 4 mM DTT.

Sgo1 fragments (Sgo1_1-415_, Sgo1_1-155_ and Sgo1_1-130_) were cloned in the pTYB11 vector (IMPACT system, New England Biolabs) which contains an Intein tag with an embedded chitin binding domain. The Intein tag is a DTT-induced self-cleavable tag that allows purification of proteins with a native N-terminus, as it leaves no extra amino acids after cleavage. Cloning of the Sgo1 in the pTYB11 vector with an N-terminal Intein-tag, allowed the purification of a Sgo1 with a native N-terminus leaving the initiator methionine exposed to be excised by Methionine Aminopeptidases (Giglione et al., 2004). Sgo1 fragments were expressed in the BL21 (*DE3*) Gold *E. coli* strain. Cells were grown at 37°C to an OD of 1.5 and induced overnight at 18° C with 0.35 mM IPTG. Cells were resuspended in lysis buffer containing 20 mM Tris-HCl pH 7.5, 500 mM NaCl, 1mM EDTA and supplemented with complete EDTA-free cocktail tablets (1 tablet/50ml cells; Roche), 0.01 mg/ml DNase (Sigma) and 1 mM PMSF. The lysate was sonicated for 8 minutes, centrifuged at 58,000 × g for 50 minutes at 4°C and the protein was batch purified using chitin beads (NEB). Protein-bound chitin beads were washed with lysis buffer and high salt buffer (20 mM Tris-HCl pH 7.5, 1 M NaCl, 1 mM EDTA and 1 mM ATP) and eluted with 20 mM Tris-HCl pH 7.5, 500 mM NaCl, 50 mM DTT overnight at room temperature. The eluted protein was then dialysed into 20 mM Tris-HCl pH 7.5, 150 mM NaCl, 50 mM Glutamate, 50 mM Arginine and 2mM DTT overnight at 4°C and loaded onto a HiTrap SP-HP (Cytiva) ion exchange column. The samples containing Sgo1 were pooled, concentrated and run in a Superdex 200 Increase 10/300 column (Cytiva) pre-equilibrated with 25 mM HEPES pH 8, 250mM NaCl, 5 % Glycerol and 2 M DTT.

Sgo1_130-280_ was cloned in a pEC-S-CDF-His vector as N-terminally His-tagged. Sgo1_130-280 4A_ was generated using the Quickchange site-directed mutagenesis method (Stratagene). The vectors were transformed in BL21 Gold cells and grown and induced as described above. Cells were resuspended in lysis buffer containing 20 mM Tris-HCl pH 8, 500 mM NaCl, 35 mM Imidazole, 2 mM β-mercaptoethanol and supplemented with complete EDTA-free cocktail tablets (1 tablet/50ml cells; Roche), 0.01 mg/ml DNase (Sigma) and 1 mM PMSF. The protein was purified using a HisTrap column (Cytiva). The protein-bound column was washed with lysis buffer and high salt buffer (20 mM Tris-HCl pH 8, 1 M NaCl, 35 mM Imidazole, 2 mM β-mercaptoethanol) and eluted with 20 mM Tris-HCl pH 8, 200 mM NaCl, 400 mM Imidazole and 2 mM β-mercaptoethanol. The eluted protein was then dialysed into 20 mM Tris-HCl pH 8, 200 mM NaCl, 1 mM DTT overnight at 4°C and loaded onto a HiTrap Q (Cytiva) ion exchange column. The samples containing Sgo1 were pooled, concentrated and run in a Superdex 200 Increase 10/300 column (Cytiva) pre-equilibrated with 25 mM HEPES pH 8, 150mM NaCl, 5 % Glycerol and 1 mM DTT.

### Interaction studies using size exclusion chromatography

All SEC experiments for the purified recombinant proteins were performed on an AKTA Pure 25 High Performance Liquid Chromatography (HPLC) unit (Cytiva) with sample collector. For all interaction studies a Superdex 200 10/300GL 24ml column (Cytiva) was used. Before sample injection, the column was pre-equilibrated in 25 mM HEPES pH 7.5, 150 mM NaCl, 4 mM DTT, 5% glycerol (v/v) for interaction experiments involving Sgo1_1-155_ or pre-equilibrated in 25 mM HEPES pH 8, 250 mM NaCl, 2 mM DTT, 5% glycerol (v/v) for interaction experiments involving Sgo1_1-415_. 0.5 ml fractions were collected with a 0.2 CV delayed fractionation setting. UV 280 nm and 260 nm wavelengths were monitored.

### Chemical cross-linking and MS analysis

Cross-linking experiments of Sgo1_1-415_ and CPC_ISB10-280_ were performed using 1-ethyl-3-(3-dimethylaminopropyl) carbodiimide (EDC, Thermo Fisher Scientific) in the presence of *N*-hydroxysulfosuccinimide (NHS, Thermo Fisher Scientific). 25 μg of gel filtrated protein complex was cross-linked with 120 μg EDC and 44 μg of NHS in 25 mM HEPES pH 6.8, 150 mM NaCl for 1h 30min at room temperature. The cross-linking was stopped by the addition of 100 mM Tris-HCl and cross-linking products were briefly resolved using a 4-12 % Bis-Tris NuPAGE (Thermo Fisher Scientific). Bands were visualised by short Instant Blue staining (Abcam), excised, reduced with 10 mM DTT for 30 min at room temperature, alkylated with 5 mM iodoacetamide for 20 min at room temperature and digested overnight at 37 °C using 13 ng/μl trypsin (Promega). Digested peptides were loaded onto C18-Stage-tips (Rappsilber et al., 2007). LC-MS/MS analysis was performed using an Orbitrap Fusion™ Lumos™ Tribrid™ Mass Spectrometer (Thermo Scientific) applying a “high-high” acquisition strategy. Peptide mixtures were injected for each mass spectrometric acquisition. Peptides were separated on a 75 µm × 50 cm PepMap EASY-Spray column (Thermo Scientific) fitted into an EASY-Spray source (Thermo Scientific), operated at 50 °C column temperature. Mobile phase A consisted of water and 0.1% v/v formic acid. Mobile phase B consisted of 80% v/v acetonitrile and 0.1% v/v formic acid. Peptides were loaded at a flow-rate of 0.3 μl/min and eluted at 0.2 μl/min using a linear gradient going from 2% mobile phase B to 40% mobile phase B over 139 (or 109) minutes, followed by a linear increase from 40% to 95% mobile phase B in 11 minutes. The eluted peptides were directly introduced into the mass spectrometer. MS data were acquired in the data-dependent mode with the top-speed option. For each three-second acquisition cycle, the mass spectrum was recorded in the Orbitrap with a resolution of 120,000. The ions with a precursor charge state between 3+ and 8+ were isolated and fragmented using HCD or EThcD. The fragmentation spectra were recorded in the Orbitrap. Dynamic exclusion was enabled with single repeat count and 60-second exclusion duration.

The mass spectrometric raw files were processed into peak lists using ProteoWizard (version 3.0.20338) (Kessner et al., 2008), and cross-linked peptides were matched to spectra using Xi software (version 1.7.6.3) (Mendes et al., 2018), https://github.com/Rappsilber-Laboratory/XiSearch) with in-search assignment of monoisotopic peaks (Lenz et al., 2018). Search parameters were MS accuracy, 3 ppm; MS/MS accuracy, 10ppm; enzyme, trypsin; cross-linker, EDC; max missed cleavages, 4; missing mono-isotopic peaks, 2; fixed modification, carbamidomethylation on cysteine; variable modifications, oxidation on methionine; fragments b and y type ions (HCD) or b, c, y, and z type ions (EThcD) with loss of H2O, NH3 and CH3SOH. 1% on link level False discovery rate (FDR) was estimated based on the number of decoy identification using XiFDR (Fischer and Rappsilber, 2017). The MS proteomics data have been deposited to the ProteomeXchange Consortium via the PRIDE (Perez-Riverol et al., 2019) partner repository.

### Isothermal Calorimetry

ITC experiments were performed using a MicroCal Auto-iTC200 (Malvern Instruments, Worcestershire, UK). A total of 40 μl of Survivin/CPC complexes 50-375 μM (monomer concentration) was injected into 200 μl of 5-20 μM hSgo1 constructs (monomer concentration) in 16 aliquots (1 × 0.5 and 15 × 2.5 μl), 180 s between injections, reference power 3 µcal.s^-1^, syringe spin 750 rpm and filter period 5s. Control titrations were performed in which the injectant was added to buffer without protein or buffer was injected into the protein. Titrations were carried out at 20 °C, except for the analysis of the Survivin/Sgo1_AKER_ interaction, that was performed at 10 °C. The heats of reaction were corrected for the heat of dilution and analysed using the MicroCal ITC Software V1.30 (Malvern Instruments). All experiments were carried out in 50 mM HEPES pH 8, 150 mM NaCl, 5 % (v/v) glycerol, 1 mM TCEP.

### Mass Photometry

High precision microscope coverslips (No. 1.5, 24 × 50 mm) were cleaned with Milli-Q water, 100 % isopropanol, Milli-Q water and dried. Silicone gaskets (Grace BioLabs 103250) were placed on the coverslips. Samples were cross-linked with 0.01% glutaraldehyde for 5 min at 4 °C and quenched by addition of 50 mM Tris-HCl pH 7.5 for 1h at 4 °C. Immediately prior to mass photometry measurements, samples were diluted to 100 nM in buffer containing 25mM HEPES pH 8, 250 mM NaCl and 2 mM DTT. For each acquisition, 20 nM of diluted protein was measured following manufacturer’s instructions. All data was acquired using a One^MP^ mass photometer instrument (Refeyn Ltd, Oxford, UK) and AcquireMP software (Refeyn Ltd., v2.4.1). Movies were recorded in the regular field of view using default settings. Data was analysed using Discover^MP^ software (Refeyn Ltd., v2.4.2).

### Tethering Assays

The LacO tethering assays were performed essentially as described before (Hadders et al., 2020). U-2 OS LacO Haspin CM cells (Hadders et al., 2020) were seeded on glass coverslips and directly transduced with recombinant baculovirus expressing LacI-GFP fusion proteins. After ∼ 4-6 hours STLC (20 μM) was added and left to incubate overnight. The next morning cells were fixed in 4 % PFA for 15 minutes and permeabilized with ice cold methanol. Prior to staining cells were blocked in PBS supplemented with 0.01 % Tween20 (PBST) and 3 % BSA for 30 minutes followed by staining with primary antibodies in PBST + 3 % BSA for 2-4 hours. Coverslips were then washed three times with PBST followed by staining with secondary antibodies and DAPI (1 μg/ml) for 1 hour. After another three washes with PBST coverslips were mounted using Prolong Diamond. Cells were imaged on a DeltaVision system. The following antibodies were used for indirect immunofluorescence: anti-Aurora B (1:1000; 611083; BD transductions), anti-Borealin (1:1000; rabbit polyclonal; a kind gift from Dr. S. Wheatley), GFP-Booster (1:1000; ABIN509419; Chromotek). The secondary antibodies used were goat anti-mouse IgG Alexa Fluor 568 conjugate (1:500; A-11031; Thermo Fisher Scientific), goat anti-mouse IgG Alexa Fluor 647 conjugate (1:500; A-1103121236; Thermo Fisher Scientific), goat anti-rabbit IgG Alexa Fluor 568 conjugate (1:500; A-11036; Thermo Fisher Scientific) and goat anti-rabbit IgG Alexa Fluor 647 conjugate (1:500; A-21245; Thermo Fisher Scientific).

### Rescue experiments and immunofluorescence microscopy

Sgo1 was cloned in the pCDNA3-GFP vector (6D) (a gift from Scott Gradia, California Institute for Quantitative Biosciences [QB3], University of California, Berkeley, CA; Addgene plasmid #30127; http://n2t.net/addgene:30127; RRID:Addgene_30127). Mutations of Sgo1 were obtained using the Quikchange site-directed mutagenesis method (Stratagene). DNA transfection (700 ng) was performed using jetPRIME (Polyplus Transfection) according to the manufacturer’s instructions. Lipofectamine RNAimax was used for depletion of endogenous Sgo1 using the following oligonucleotide: 5′-UGCACCAUGCCAAUAAdTdT-3′ (40 pmol). Luciferase targeting was used as a control (5′-CGUACGCGGAAUACUUCGAdTdT-3′; Elbashir et al., 2001). All siRNA oligonucleotides were purchased from Qiagen. Cells were plated in glass coverslips, transfected with jetPRIME (DNA) 16 h after plating and transfected with Lipofectamine siRNA max (siRNA oligos) 24 h after the first transfection. For centromeric quantification of the Borealin signal, HeLa Kyoto cells were synchronized with 50 ng/ml nocodazole for 16 h, 8 h after siRNA transfection. A minimum of 50 cells per condition were quantified. The acquired images were processed by constrained iterative deconvolution using SoftWoRx 3.6 software package (Applied Precision), and the centromere intensity of Borealin was quantified using an ImageJ plugin (https://doi.org/10.5281/zenodo.5145584). Briefly, the plugin quantifies the mean fluorescence signal of Borealin in a 3-pixel-wide ring immediately outside the centromere, defined by the ACA staining. For background subtraction, a selected area within the cytoplasm signal was selected. To compare data from different replicates, values obtained after background correction were averaged and normalized to the mean of Borealin intensity in the Sgo1-GFP rescue condition. Statistical significance of the difference between normalized intensities at the centromere region was established by a Mann–Whitney *U* test using Prism 6.0.

Quantification of anaphases displaying chromosome bridges or lagging chromosomes was performed 24 h after HeLa Kyoto cells were transfected with the siRNA oligos. For the Monastrol assay, HeLa Kyoto cells were synchronised with 100 μM Monastrol for 16 h and released into 5 μM MG132 for 1 h. The experiments were performed in triplicate and a minimum of 85 cells per condition were quantified.

For the chromosome spreads, 8 h after siRNA oligo transfection, HeLa cells were treated with 50 ng/ml Nocodazole. 16 h after Nocodazole treatment, cells were collected by mitotic shake off and incubated in hypotonic buffer (75 mM KCl) at 37 °C for 10 min. After attachment to glass coverslips using Cytospin at 1,800 rpm for 5 min, chromosome spreads were extracted with ice-cold PBS-0.2 % triton for 4 min and fixed with 4 % PFA. The immunofluorescence was performed as described below. Three replicates were performed and a minimum of 49 kinetochores were analysed. The centroids of kinetochores were detected in ImageJ using Speckle TrackerJ (Smith et al., 2011) software. A custom ImageJ script (DOI: 10.5281/zenodo.5235670) was then used to assign kinetochore pairs as closest neighbours, with a maximum separation of 1.5µm. The fluorescence intensities along 2 µm line ROIs through the centroids and centred on the midpoint of the pair were taken in both the channels. Full width half max values for the Borealin line plots were calculated by linear interpolation using a combination of the point-slope formula and the slope formula. Statistical significance of the difference between the full width half max values between different Sgo1 constructs was established by a Mann–Whitney *U* test using Prism 6.0.

In all cases, cells were fixed in 4% paraformaldehyde (PFA) 48 h after DNA transfection and 24 h after oligonucleotide transfection. Cells were then permeabilised with permeabilization buffer (0.2 % Triton in 1X PBS) for 10 min, blocked with 3 % BSA in permeabilization buffer for 1 h and incubated with primary and secondary antibodies in blocking buffer for 1 h each. All experiments were performed in triplicate.

The following antibodies were used for indirect immunofluorescence: anti-Sgo1 (1:300; kind gift from Ana Losada, Spanish National Cancer Research Centre (CNIO), Madrid, Spain), anti-Borealin (1:500; 147-3; MBL), anti-tubulin (1:2000; B512; Sigma), anti-CENP-C (1:400; kind gift from William C. Earnshaw, Wellcome Centre for Cell Biology, University of Edinburgh, Edinburgh, UK) and anti-ACA (1:300; 15-235; Antibodies Inc.). The secondary antibodies used were FITC-conjugated AffiniPure goat anti-rabbit IgG, TRITC-conjugated AffiniPure goat anti-rabbit IgG, TRITC-conjugated AffiniPure donkey anti-mouse, Cy5-conjugated AffiniPure donkey anti-human, Cy5-conjugated AffiniPure donkey anti-mouse (1:300; Jackson Immunoresearch). DAPI was used for DNA staining. Imaging was performed at room temperature using a wide-field DeltaVision Elite (Applied Precision) microscope with Photometrics Cool Snap HP camera and 100× NA 1.4 Plan Apochromat objective with oil immersion (refractive index = 1.514) using the SoftWoRx 3.6 (Applied Precision) software. Shown images are deconvolved and maximum-intensity projections.

### Western blot

To study the Sgo1 levels after siRNA oligo treatment and to test the expression levels of each of the Sgo1-GFP constructs, HeLa Kyoto cells were transfected in 12-well dishes as described above and lysed in 1× Laemmli buffer, boiled for 5 min, and analysed by SDS-PAGE followed by Western blotting. The antibodies used for the immunoblot were anti-Sgo1 antibody [1:1000; a gift from Ana Losada’s laboratory, Spanish National Cancer Research Centre (CNIO), Madrid, Spain; Serrano et al., 2009], anti-tubulin (1:10,000; ab18251; Abcam) and anti-GFP (1:1000; Abcam). Secondary antibodies used were goat anti-mouse 680, donkey anti-rabbit 800 and donkey anti-mouse 800 (LI-COR) at 1:2,000 dilution. Immunoblots were imaged using the Odyssey CLx system.

## Acknowledgments

We thank the staff of the Edinburgh Protein Production Facility and the Centre for Optical Instrumentation Laboratory for their help. We also thank Ana Losada (Chromosome Dynamics Group, Molecular Oncology Programme, Spanish National Cancer Research Centre [CNIO], Madrid, Spain) for kindly providing the Sgo1 antibody and William C. Earnshaw (Wellcome Centre for Cell Biology, University of Edinburgh, Edinburgh, UK) for kindly providing the CENP-C antibody. We also thank Refeyn Ltd (Oxford, UK), specially Akhila Bettadapur and Tomás de Garay for their help with the mass photometry experiments. We also thank Cristina Ferrás and Marco Cruz (Instituto de Biologia Molecular e Celular, Universidade do Porto, Portugal) for sharing with us the Sgo1 siRNA oligo sequence. We thank Bethan Medina-Pritchard for critical reading of the manuscript.

The Wellcome Trust generously supported this work through a Career Development and Enhancement Grants (095822) and Senior Research Fellowship (202811) to A.A. Jeyaprakash, a Centre Core Grant (203149) and a Core Grant (109916/Z/15/Z) to the Edinburgh Protein Production Facility. This study was also supported by a research grant from the Dutch Cancer Society (KWF grant 10366) to S.M.A Lens. Moreover, the Lens lab is part of Oncode Institute which is partly financed by the Dutch Cancer Society. Tanmay Gupta was funded by the Darwin Trust of Edinburgh.

The authors declare no competing financial interests.

## Author contributions

A.A. Jeyaprakash conceived the project. M.A. Abad, T. Gupta, M. A. Hadders, A. Meppelink, J. P. Wopken, E. Blackburn, D. Kelly, T. McHugh, J. Rappsilber and S. M. A. Lens designed the experiments. M.A. Abad, T. Gupta, E. Blackburn, J. Zou and L. Buzuk performed the biochemical, structural and biophysical characterisation. M. A. Hadders, A. Meppelink and J. P. Wopken performed the tethering assays. M.A. Abad performed the functional assays. M.A. Abad, M. A. Hadders, E. Blackburn, S. M. A. Lens and A.A. Jeyaprakash wrote the manuscript.

## References

Bekier, M.E., T. Mazur, M.S. Rashid, and W.R. Taylor. 2015. Borealin dimerization mediates optimal CPC checkpoint function by enhancing localization to centromeres and kinetochores. Nature Communications. 6:6775.

Bonner, M.K., J. Haase, H. Saunders, H. Gupta, B.I. Li, and A.E. Kelly. 2020. The Borealin dimerization domain interacts with Sgo1 to drive Aurora B–mediated spindle assembly. Molecular Biology of the Cell. 31:2207–2218.

Bourhis, E., A. Lingel, Q. Phung, W.J. Fairbrother, and A.G. Cochran. 2009. Phosphorylation of a borealin dimerization domain is required for proper chromosome segregation. Biochemistry. 48:6783–6793.

Broad, A.J., K.F. DeLuca, and J.G. DeLuca. 2020. Aurora B kinase is recruited to multiple discrete kinetochore and centromere regions in human cells. J Cell Biol. 219.

Carmena, M., M. Wheelock, H. Funabiki, and W.C. Earnshaw. 2012. The chromosomal passenger complex (CPC): from easy rider to the godfather of mitosis. Nature Reviews Molecular Cell Biology. 13:789.

Cheeseman, I.M., J.S. Chappie, E.M. Wilson-Kubalek, and A. Desai. 2006. The conserved KMN network constitutes the core microtubule-binding site of the kinetochore. Cell. 127:983–997.

Cimini, D., X. Wan, C.B. Hirel, and E.D. Salmon. 2006. Aurora kinase promotes turnover of kinetochore microtubules to reduce chromosome segregation errors. Curr Biol. 16:1711–1718.

DeLuca, J.G., W.E. Gall, C. Ciferri, D. Cimini, A. Musacchio, and E.D. Salmon. 2006. Kinetochore microtubule dynamics and attachment stability are regulated by Hec1. Cell. 127:969–982.

Du, J., A.E. Kelly, H. Funabiki, and D.J. Patel. 2012. Structural basis for recognition of H3T3ph and Smac/DIABLO N-terminal peptides by human Survivin. Structure (London, England : 1993). 20:185–195.

Fischer, L., and J. Rappsilber. 2017. Quirks of Error Estimation in Cross-Linking/Mass Spectrometry. Analytical chemistry. 89:3829–3833.

Foley, E.A., and T.M. Kapoor. 2013. Microtubule attachment and spindle assembly checkpoint signalling at the kinetochore. Nat Rev Mol Cell Biol. 14:25–37.

Funabiki, H., and D.J. Wynne. 2013. Making an effective switch at the kinetochore by phosphorylation and dephosphorylation. Chromosoma. 122:135–158.

Gandhi, R., P.J. Gillespie, and T. Hirano. 2006. Human Wapl is a cohesin-binding protein that promotes sister-chromatid resolution in mitotic prophase. Curr Biol. 16:2406–2417.

Gelens, L., J. Qian, M. Bollen, and A.T. Saurin. 2018. The Importance of Kinase–Phosphatase Integration: Lessons from Mitosis. Trends in Cell Biology. 28:6–21.

Giglione, C., A. Boularot, and T. Meinnel. 2004. Protein N-terminal methionine excision. Cell Mol Life Sci. 61:1455–1474.

Hadders, M.A., S. Hindriksen, M.A. Truong, A.N. Mhaskar, J.P. Wopken, M.J.M. Vromans, and S.M.A. Lens. 2020. Untangling the contribution of Haspin and Bub1 to Aurora B function during mitosis. J Cell Biol. 219.

Haering, C.H., A.M. Farcas, P. Arumugam, J. Metson, and K. Nasmyth. 2008. The cohesin ring concatenates sister DNA molecules. Nature. 454:297–301.

Haering, C.H., J. Löwe, A. Hochwagen, and K. Nasmyth. 2002. Molecular architecture of SMC proteins and the yeast cohesin complex. Mol Cell. 9:773–788.

Hengeveld, R.C.C., M.J.M. Vromans, M. Vleugel, M.A. Hadders, and S.M.A. Lens. 2017. Inner centromere localization of the CPC maintains centromere cohesion and allows mitotic checkpoint silencing. Nat Commun. 8:15542.

Hindriksen, S., S.M.A. Lens, and M.A. Hadders. 2017. The Ins and Outs of Aurora B Inner Centromere Localization. Frontiers in cell and developmental biology. 5:112.

Janicki, S.M., T. Tsukamoto, S.E. Salghetti, W.P. Tansey, R. Sachidanandam, K.V. Prasanth, T. Ried, Y. Shav-Tal, E. Bertrand, R.H. Singer, and D.L. Spector. 2004. From silencing to gene expression: real-time analysis in single cells. Cell. 116:683–698.

Jeyaprakash, A.A., C. Basquin, U. Jayachandran, and E. Conti. 2011. Structural basis for the recognition of phosphorylated histone h3 by the survivin subunit of the chromosomal passenger complex. Structure. 19:1625–1634.

Jeyaprakash, A.A., U.R. Klein, D. Lindner, J. Ebert, E.A. Nigg, and E. Conti. 2007. Structure of a Survivin–Borealin–INCENP Core Complex Reveals How Chromosomal Passengers Travel Together. Cell. 131:271–285.

Kawashima, S.A., T. Tsukahara, M. Langegger, S. Hauf, T.S. Kitajima, and Y. Watanabe. 2007. Shugoshin enables tension-generating attachment of kinetochores by loading Aurora to centromeres. Genes Dev. 21:420–435.

Kawashima, S.A., Y. Yamagishi, T. Honda, K. Ishiguro, and Y. Watanabe. 2010. Phosphorylation of H2A by Bub1 prevents chromosomal instability through localizing shugoshin. Science. 327:172–177.

Kelly, A.E., C. Ghenoiu, J.Z. Xue, C. Zierhut, H. Kimura, and H. Funabiki. 2010. Survivin Reads Phosphorylated Histone H3 Threonine 3 to Activate the Mitotic Kinase Aurora B. Science. 330:235LP–239.

Kessner, D., M. Chambers, R. Burke, D. Agus, and P. Mallick. 2008. ProteoWizard: open source software for rapid proteomics tools development. Bioinformatics (Oxford, England). 24:2534–2536.

Kitajima, T.S., T. Sakuno, K.-i. Ishiguro, S.-i. Iemura, T. Natsume, S.A. Kawashima, and Y. Watanabe. 2006. Shugoshin collaborates with protein phosphatase 2A to protect cohesin. Nature. 441:46–52.

Kolbowski, L., C. Combe, and J. Rappsilber. 2018. xiSPEC: web-based visualization, analysis and sharing of proteomics data. Nucleic acids research. 46:W473–W478.

Kueng, S., B. Hegemann, B.H. Peters, J.J. Lipp, A. Schleiffer, K. Mechtler, and J.M. Peters. 2006. Wapl controls the dynamic association of cohesin with chromatin. Cell. 127:955–967.

Lampson, M.A., K. Renduchitala, A. Khodjakov, and T.M. Kapoor. 2004. Correcting improper chromosome-spindle attachments during cell division. Nat Cell Biol. 6:232–237.

Lenz, S., S.H. Giese, L. Fischer, and J. Rappsilber. 2018. In-Search Assignment of Monoisotopic Peaks Improves the Identification of Cross-Linked Peptides. Journal of proteome research. 17:3923–3931.

Liang, C., Z. Zhang, Q. Chen, H. Yan, M. Zhang, L. Zhou, J. Xu, W. Lu, and F. Wang. 2019. Centromere-localized Aurora B kinase is required for the fidelity of chromosome segregation. Journal of Cell Biology. 219.

Liu, H., L. Jia, and H. Yu. 2013a. Phospho-H2A and Cohesin Specify Distinct Tension-Regulated Sgo1 Pools at Kinetochores and Inner Centromeres. Current Biology. 23:1927–1933.

Liu, H., S. Rankin, and H. Yu. 2013b. Phosphorylation-enabled binding of SGO1-PP2A to cohesin protects sororin and centromeric cohesion during mitosis. Nature cell biology. 15:40–49.

Mendes, M.L., L. Fischer, Z.A. Chen, M. Barbon, F.J. O’Reilly, M. Bohlke-Schneider, A. Belsom, T. Dau, C.W. Combe, M. Graham, M.R. Eisele, W. Baumeister, C. Speck, and J. Rappsilber. 2018. An integrated workflow for cross-linking/mass spectrometry. bioRxiv:355396.

Meppelink, A., L. Kabeche, Martijn J.M. Vromans, Duane A. Compton, and Susanne M.A. Lens. 2015. Shugoshin-1 Balances Aurora B Kinase Activity via PP2A to Promote Chromosome Bi-orientation. Cell Reports. 11:508–515.

Musacchio, A. 2015. The Molecular Biology of Spindle Assembly Checkpoint Signaling Dynamics. Current Biology. 25:R1002–R1018.

Musacchio, A., and A. Desai. 2017. A Molecular View of Kinetochore Assembly and Function. Biology (Basel). 6.

Niedzialkowska, E., F. Wang, P.J. Porebski, W. Minor, J.M. Higgins, and P.T. Stukenberg. 2012. Molecular basis for phosphospecific recognition of histone H3 tails by Survivin paralogues at inner centromeres. Mol Biol Cell. 23:1457–1466.

Peplowska, K., A.U. Wallek, and Z. Storchova. 2014. Sgo1 Regulates Both Condensin and Ipl1/Aurora B to Promote Chromosome Biorientation. PLOS Genetics. 10:e1004411.

Salic, A., J.C. Waters, and T.J. Mitchison. 2004. Vertebrate shugoshin links sister centromere cohesion and kinetochore microtubule stability in mitosis. Cell. 118:567–578.

Saurin, A.T. 2018. Kinase and Phosphatase Cross-Talk at the Kinetochore. Frontiers in Cell and Developmental Biology. 6.

Serrano, A., M. Rodríguez-Corsino, and A. Losada. 2009. Heterochromatin protein 1 (HP1) proteins do not drive pericentromeric cohesin enrichment in human cells. PLoS One. 4:e5118.

Shintomi, K., and T. Hirano. 2009. Releasing cohesin from chromosome arms in early mitosis: opposing actions of Wapl-Pds5 and Sgo1. Genes Dev. 23:2224–2236.

Smith, Matthew B., E. Karatekin, A. Gohlke, H. Mizuno, N. Watanabe, and D. Vavylonis. 2011. Interactive, Computer-Assisted Tracking of Speckle Trajectories in Fluorescence Microscopy: Application to Actin Polymerization and Membrane Fusion. Biophysical Journal. 101:1794–1804.

Tsukahara, T., Y. Tanno, and Y. Watanabe. 2010. Phosphorylation of the CPC by Cdk1 promotes chromosome bi-orientation. Nature. 467:719.

van der Waal, M.S., A.T. Saurin, M.J. Vromans, M. Vleugel, C. Wurzenberger, D.W. Gerlich, R.H. Medema, G.J. Kops, and S.M. Lens. 2012. Mps1 promotes rapid centromere accumulation of Aurora B. EMBO Rep. 13:847–854.

Waizenegger, I.C., S. Hauf, A. Meinke, and J.M. Peters. 2000. Two distinct pathways remove mammalian cohesin from chromosome arms in prophase and from centromeres in anaphase. Cell. 103:399–410.

Wang, F., J. Dai, J.R. Daum, E. Niedzialkowska, B. Banerjee, P.T. Stukenberg, G.J. Gorbsky, and J.M.G. Higgins. 2010. Histone H3 Thr-3 Phosphorylation by Haspin Positions Aurora B at Centromeres in Mitosis. Science. 330:231LP–235.

Waterhouse, A.M., J.B. Procter, D.M.A. Martin, M. Clamp, and G.J. Barton. 2009. Jalview Version 2--a multiple sequence alignment editor and analysis workbench. Bioinformatics (Oxford, England). 25:1189–1191.

Welburn, J.P., M. Vleugel, D. Liu, J.R. Yates, 3rd, M.A. Lampson, T. Fukagawa, and I.M. Cheeseman. 2010. Aurora B phosphorylates spatially distinct targets to differentially regulate the kinetochore-microtubule interface. Mol Cell. 38:383–392.

Yamagishi, Y., T. Honda, Y. Tanno, and Y. Watanabe. 2010. Two Histone Marks Establish the Inner Centromere and Chromosome Bi-Orientation. Science. 330:239LP–243.

