## Supplementary Figures for "Molecular Basis for CPC-Sgo1 Interaction: Implications for Centromere Localisation and Function of the CPC"

### Figure S1

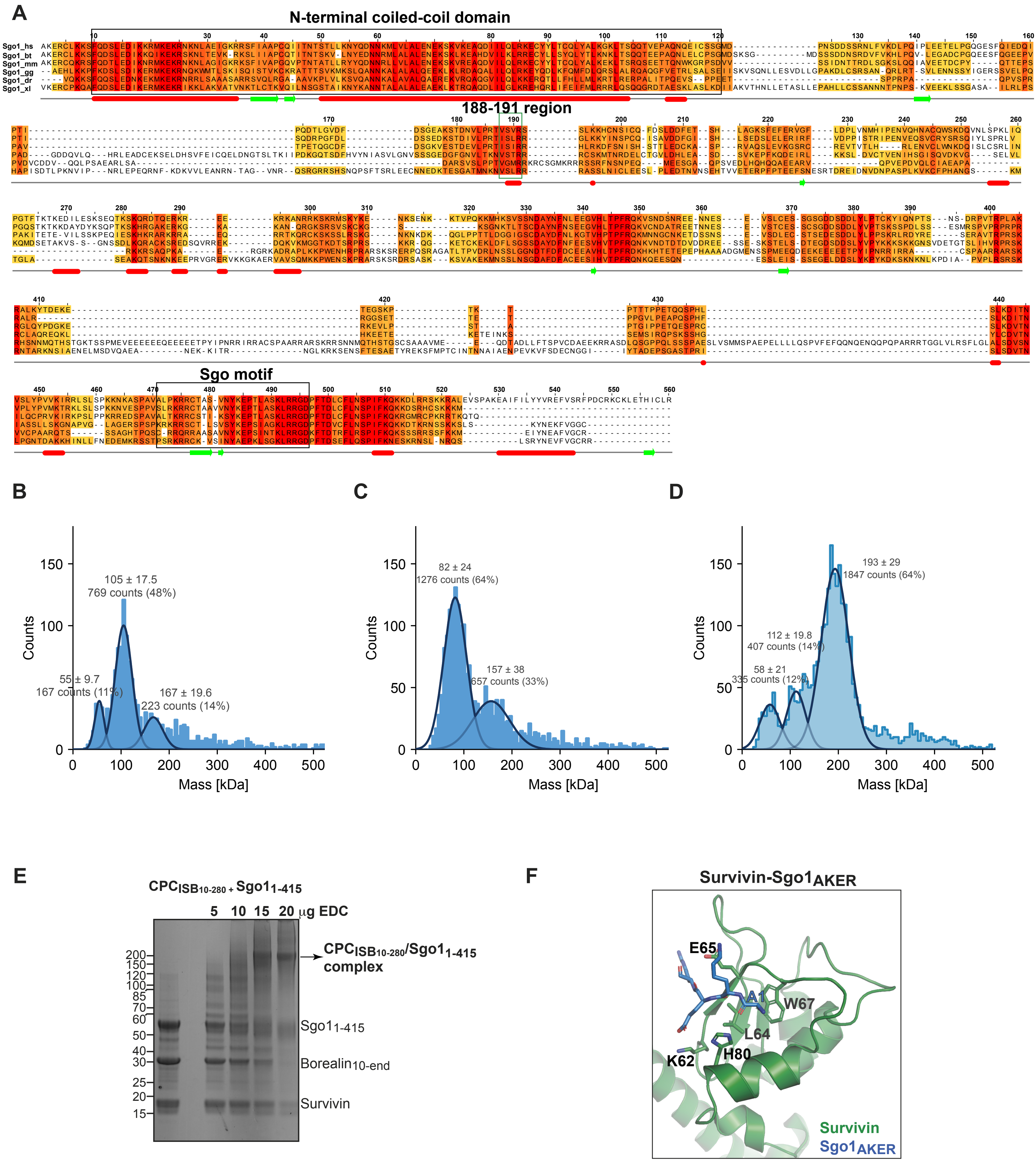
A
N-terminal coiled-coil domain
188-191 region
Sgo motif
Sgo1_hs
Sgo1_bt
Sgo1_mm
Sgo1_gg
Sgo1_dr
Sgo1_xl
B
105 ± 17.5
769 counts (48%)
55 ± 9.7
167 counts (11%)
167 ± 19.6
223 counts (14%)
Counts
Mass [kDa]
C
82 ± 24
1276 counts (64%)
157 ± 38
657 counts (33%)
Counts
Mass [kDa]
D
193 ± 29
1847 counts (64%)
112 ± 19.8
407 counts (14%)
58 ± 21
335 counts (12%)
Counts
Mass [kDa]
E
CPCISB10-280 + Sgo11-415
5 10 15 20 μg EDC
CPCISB10-280/Sgo11-415 complex
Sgo11-415
Borealin10-end
Survivin
F
Survivin-Sgo1AKER
E65
A1
W67
L64
K62
H80
Survivin
Sgo1AKER

### Figure S2

A

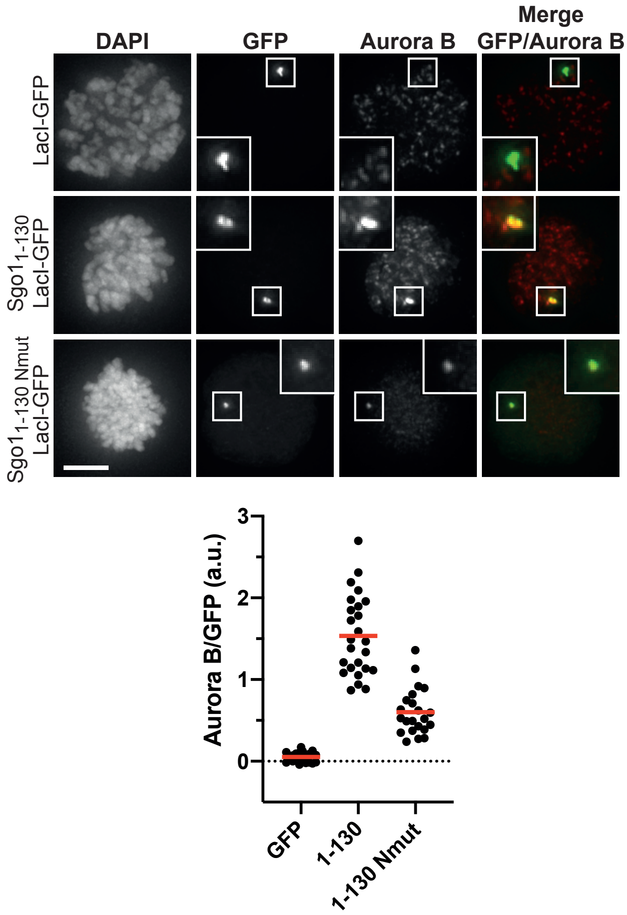


B

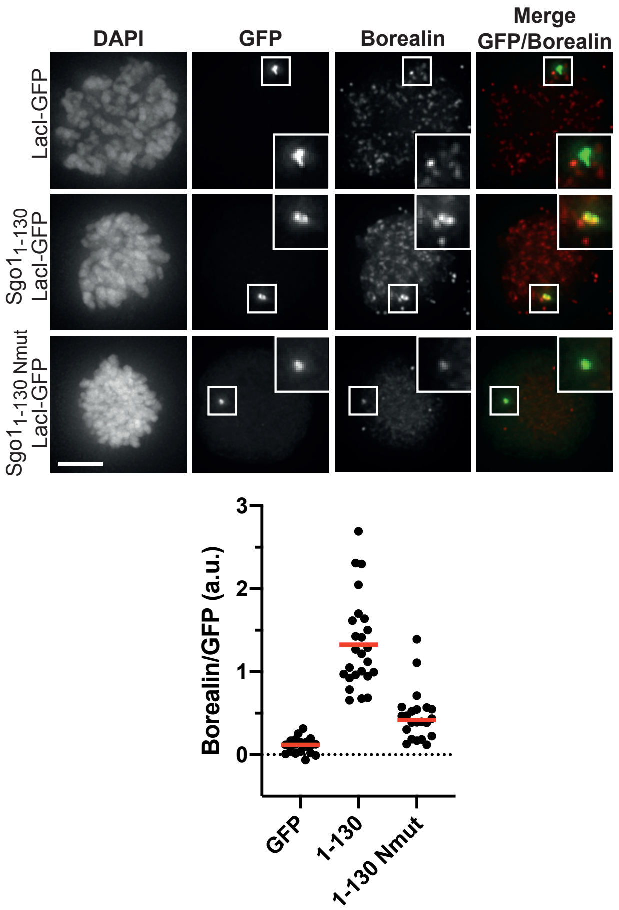


C

| Protein | N | $K_d$ (nM) | ΔG (kcal.mol$^{-1}$) | ΔH (kcal.mol$^{-1}$) | -TΔS (kcal.mol$^{-1}$K$^{-1}$) |
|---|---|---|---|---|---|
| $CPC_{ISB10\text{-}280}$ + $Sgo1_{1\text{-}415}$ | 0.726 ± 0.0057 | 52.8 ± 6.95 | -9.77 | -6.58 ± 0.098 | -3.19 |
| Survivin + $Sgo1_{1\text{-}415}$ | 0.966 ± 0.0072 | 255 ± 33 | -8.84 | -5.65 ± 0.076 | -3.20 |
| $CPC_{ISB10\text{-}280}$ + $Sgo1_{1\text{-}130}$ | 0.866 ± 0.0049 | 57.4 ± 7.92 | -9.72 | -6.22 ± 0.083 | -3.49 |
| $CPC_{ISB10\text{-}221}$ + $Sgo1_{1\text{-}130}$ | 0.903 ± 0.0043 | 163 ± 15.9 | -9.11 | -4.67 ± 0.044 | -4.44 |
| Survivin + AKER peptide | 0.823 ± 0.0054 | 11600 ± 2220 | -6.40 | -7.83 ± 1.03 | 1.43 |

D

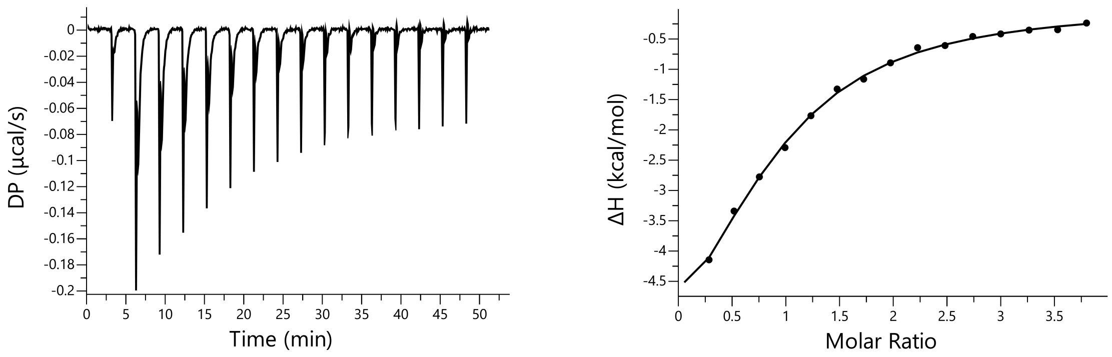

### Figure S3

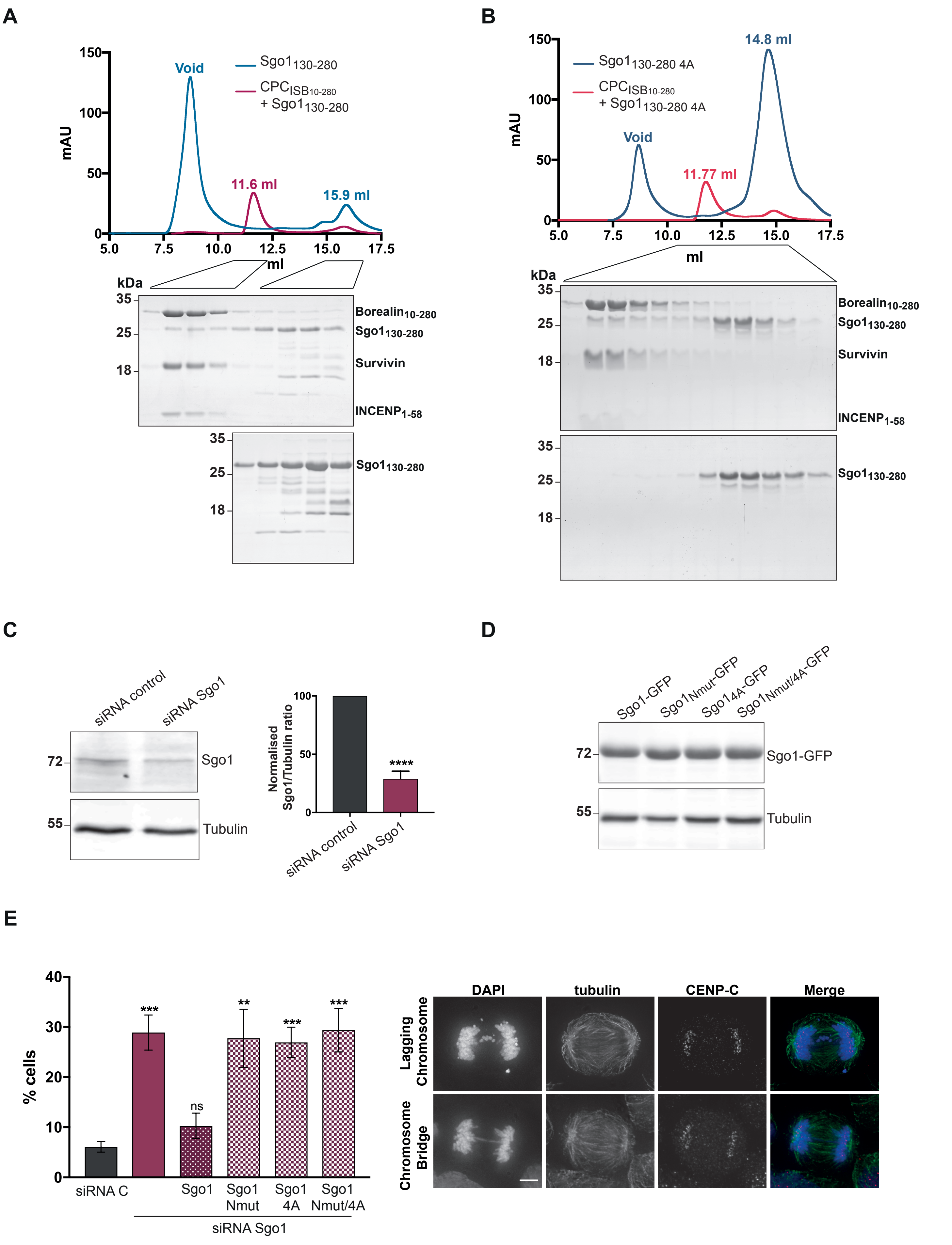
A
B
C
D
E
Void
11.6 ml
15.9 ml
Sgo1130-280
CPCISB10-280
+ Sgo1130-280
mAU
ml
kDa
Borealin10-280
Survivin
INCENP1-58
14.8 ml
11.77 ml
Sgo1130-280 4A
+ Sgo1130-280 4A
siRNA control
siRNA Sgo1
Sgo1
Tubulin
Normalised Sgo1/Tubulin ratio
****
Sgo1-GFP
Sgo1Nmut-GFP
Sgo14A-GFP
Sgo1Nmut/4A-GFP
% cells
siRNA C
Sgo1 Nmut
Sgo1 4A
Sgo1 Nmut/4A
ns
DAPI
tubulin
CENP-C
Merge
Lagging Chromosome
Chromosome Bridge
